# HAMdetector: A Bayesian regression model that integrates information to detect HLA-associated mutations

**DOI:** 10.1101/2021.07.17.452375

**Authors:** Daniel Habermann, Hadi Kharimzadeh, Andreas Walker, Yang Li, Rongge Yang, Zabrina L. Brumme, Jörg Timm, Michael Roggendorf, Daniel Hoffmann

## Abstract

**Motivation:** A key process in anti-viral adaptive immunity is that the Human Leukocyte Antigen system (HLA) presents epitopes as Major Histocompatibility Complex I (MHC I) protein-peptide complexes on cell surfaces and in this way alerts CD8^+^ cytotoxic T-Lymphocytes (CTLs). This pathway exerts strong selection pressure on viruses, favoring viral mutants that escape recognition by the HLA/CTL system, e.g. by point mutations that decrease binding of viral peptides to MHC I. Naturally, such immune escape mutations often emerge in highly variable viruses, e.g. HIV or HBV, as HLA-associated mutations (HAMs), specific to the host HLA alleles and its MHC I proteins. The reliable identification of HAMs is not only important for understanding viral genomes and their evolution, but it also impacts the development of broadly effective anti-viral treatments and vaccines against variable viruses.

By their very nature HAMs are amenable to detection by statistical methods in paired sequence / HLA data. However, HLA alleles are very polymorphic in the human host population which makes the available data relatively sparse and noisy. Under these circumstances, one way to optimize HAM detection is to integrate all relevant information in a coherent model. Bayesian inference offers a principled approach to achieve this.

**Results:** We present a new regression model for the detection of HAMs. As we choose a Bayesian approach we can include the novel sparsity-inducing priors, and we obtain easily interpretable quantitative information on HAM candidates. The basic model can be extended to include prior information relevant to HAM detection, which we demonstrate by integrating predictions of epitope affinities to MHC I, predictions of epitope peptide processing, and computation of phylogenetic background. This integrative method improves performance in HAM detection considerably over state-of-the-art methods.

**Availability:** The source code of this software is available at https://github.com/HAMdetector/Escape.jl under a permissive MIT license.

## 1. Introduction

### 1.1 The HLA system

The human immune system recognizes viral infections through two pathways: The innate and adaptive immune response. T-cell, or “cellular”, immunity, which represents one major arm of the adaptive immune system, is modulated by Human Leukocyte Antigen (HLA) molecules (Germain, 1994): Briefly, proteins that are synthesized within the cell –which will include viral proteins if the cell is infected–, are degraded in proteasomes to peptides (Goldberg et al., 2002). Some of these peptides are presented as epitopes on the cell surface by HLA class I molecules. These viral peptide-HLA complexes can then be recognized by circulating CD8^+^ Cytotoxic T-Lymphocytes (CTLS) through their T-cell receptor (Murata et al., 2007). Following this recognition, the CTL can eliminate the infected cell (Harty et al., 2000).

HLA class I molecules are encoded at three loci, HLA-A, -B and -C, and these genes are very polymorphic with more than 20,000 known alleles in humans (Robinson et al., 2014). HLA molecules vary drastically in their affinities to given epitopes so that cells from different individuals, in general, present different peptides on the cell surface. In other words, the HLA class I alleles expressed by a given individual will determine their CTL response to a given viral pathogen.

### 1.2 HLA escape

Virus variants arise continuously through mutation. Because the HLA system modulates CTL responses through viral epitope presentation, it exerts strong selection pressure towards virus variants that escape CTL recognition (Borrow et al., 1997). Such variants could, for example, carry mutations that reduce binding of viral epitopes to HLA, or that reduce recognition of the epitope/HLA complex by the CTL’s T cell receptor, or that alter peptide processing so that epitopes are no longer presented on the infected cell surface (Yewdell and Hill, 2002).

HLA diversity drives viral evolution in individuals where a virus adapts to the specific HLA alleles expressed in the host, and in human populations, where circulating viruses adapt to HLA alleles commonly expressed in that population (Kawashima et al., 2009). Upon transmission to a new host with different HLA alleles, HLA escape mutations may revert, particularly if they are associated with a reduction in viral replication capacity Matthews et al. (2008), but they can also persist, leading to their population-level accumulation (Kawashima et al., 2009).

Whether and how quickly a given escape mutation is selected in a host depends, e.g., on the viral genomic background, the magnitude of the reduction in viral replication caused by changes in the viral proteins, the selection of compensatory mutations that recover fitness, and the strength of immune response targeting the presented epitope (Kløverpris et al., 2016).

Immune escape is a driver of viral evolution in individuals and populations, particularly for highly variable viruses such as HIV or HBV (Alizon et al., 2011; Allen et al., 2005; Rousseau et al., 2008; Lumley et al., 2018). Methods to accurately detect immune escape mutations are therefore critical. More broadly, an improved understanding of immune escape can aid in the development of treatments and vaccines that rely on effective immune responses.

### 1.3 Identifying HLA escape mutations

There are several experimental methods available to study HLA escape (Czerkinsky et al., 1983; Brunner et al., 1968; Lamoreaux et al., 2006; Altman et al., 1996). However, these methods are relatively slow and costly, especially for screening purposes. A promising approach that makes efficient use of frequently available data is to combine viral genome sequencing, host HLA determination, computational identification by statistical association analysis, and targeted experimental validation (Carlson et al., 2012).

As the selection pressure exerted by cytotoxic T cells depends on successful recognition of viral peptides bound to HLA molecules on the infected cell surface, escape mutations are HLA allele specific and can therefore be detected as HLA allele dependent amino acid substitutions, or “footprints,” in sequence alignments of viral proteins (Moore, 2002). Amino acid substitutions enriched in viral sequences from hosts with a specific HLA allele are termed HLA associated mutations (HAM).

One way of quantifying this enrichment is Fisher’s exact test (Fisher, 1922): For a given substitution *S*_*i*_ at alignment position *i* and HLA allele *H*, a 2-by-2 contingency table is constructed containing the absolute counts of the number of sequences in the four possible categories (*S*_*i*_, *H*), (*S*_*i*_, ¬*H*), (¬*S*_*i*_, *H*) and (¬*S*_*i*_, ¬*H*), where ¬*S*_*i*_ denotes any substitution except *S*_*i*_, and ¬*H* denotes any HLA allele except *H*.

Fisher’s exact test is a conventional null hypothesis significance test (NHST) that generates p-values. In this case, the null hypothesis is that HLA allele *H* and substitution *S*_*i*_ are independent, and the p-value is the probability of observing a deviation from independence that is at least as extreme as in the data at hand under the assumption that the null hypothesis is true.

Fisher’s exact test has the advantage of being fast and easy to apply (Budeus et al., 2016), but it also has several disadvantages (Carlson et al., 2008). The most striking one is that viral sequences share a common phylogenetic history, and, therefore, treating sequences as independent and identically distributed samples may under- or overestimate effect sizes. In the context of hypothesis testing, this leads to increased false positive and false negative rates (Osborne and Waters, 2002; Scariano and Davenport, 1987).

Another issue with Fisher’s exact test is the genomic proximity of human HLA class I loci (Francke and Pellegrino, 1977) leading to linkage disequilibrium – inheritance of HLA alleles can be correlated. Therefore, spurious HAMs can occur if associations of substitutions with individual HLA alleles are tested: if HLA allele *H*_1_ is associated with an amino acid substitution *R* because of immune escape, but *H*_1_ is in linkage disequilibrium with allele *H*_2_, then this leads to an association of *R* and *H*_2_, even without being an escape mutation from *H*_2_.

Carlson et al. (2008) developed the Phylogenetic Dependency Network, a method that accounts for several of the aforementioned problems, in particular phylogenetic bias and HLA linkage disequilibrium. However, it is based on null hypothesis significance testing.

### 1.4 Issues with p-values for screening

There are fundamental statistical issues with p-values as a screening tool (Amrhein and Greenland, 2017): with small effect sizes and high variance between measurements, as is often the case with biological data, statistically significant results can be misleading, can have the wrong direction (type S error), or can greatly overestimate an effect (type M error) (Gelman and Carlin, 2014). Such problems are more and more appreciated in the context of the current “replication crisis” – in the life sciences scientific claims with seemingly strong statistical support often fail to replicate (Ioannidis, 2005; Begley and Ellis, 2012; Baker, 2016).

These problems are exacerbated if p-values are used for screening purposes (multiple testing problem). The probability of obtaining a statistically significant result increases with each additional test, even in absence of any real effect. When using p-values as a filter, it is therefore likely to obtain significant effects that are in fact not real. A common strategy to mitigate this problem is to control the false discovery rate (Benjamini and Hochberg, 1995). The downside of such adjustment procedures is that only the very largest effects remain if large datasets are screened.

Instead of performing many hypothesis tests and trying to adjust for them, we prefer to fit a single, multilevel model that contains all comparisons of interest. Multilevel models can make the problem of multiple comparisons disappear entirely and yield more valid estimates (Gelman et al., 2012).

## 2. Materials and Methods

Our general approach for HAMdetector is to fit Bayesian regression models that captures relationships between host HLA alleles and substitutions in viral proteomes.

This Bayesian approach is advantageous because it allows use of: (1) prior information (e.g. knowledge of effect magnitudes), (2) relevant additional information (phylogeny, epitope information), (3) a problem-specific structure, (4) partial pooling (Gelman, 2010).

### 2.1 Model backbone

We chose a logistic regression model as backbone because it is easily extensible, and because coefficients can be interpreted in the familiar way as summands on the log-odds scale.

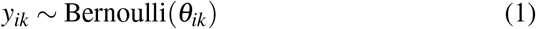

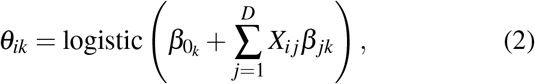

where *y*_*ik*_ is the binary encoded observation of substitution *k* in viral sequence *i* (each observed amino acid state *k* contributes a separate column to *y*_*ik*_); *θ*_*ik*_ is the estimated probability that we observe substitution *k* in sequence *i*; *β*_0*k*_ is an intercept for substitution *k*, corresponding to the overall log-odds for substitution *k*; *X*_*i j*_ is 1 if sequence *i* comes from host individual with HLA allele *j* and 0 otherwise; *β*_*jk*_ is the HLA regression coefficient of HLA allele *j* for substitution *k*; *D* is the number of HLA alleles in the dataset; the logistic inverse link function transforms the linear model in parentheses to the probability scale of *θ*_*ik*_.

The main parameters of interest for HAMdetector are the regression coefficients *β*_*jk*_, as they quantify the strength of association between the occurrence of substitution *k* and each of the observed HLA alleles. The *β*_*jk*_ are on the log-odds scale, i.e., if we go from viral sequences from hosts without HLA allele *j* to those from hosts with *j*, the log-odds log(*p*_*k*_*/*(1*− p*_*k*_)) of observing substitution *k* increase by addition of *β*_*jk*_.

Reasoning about coefficients on the log-odds scale can sometimes be unintuitive. A useful approximation to interpret logistic regression coefficients on the probability scale is the so-called divide-by-4 rule, which means that a regression coefficient of 2 corresponds to an expected increase on the probability scale of up to 2/4 = 50%.

### 2.2 Inclusion of additional information

On top of the paired data of viral sequences and host HLA alleles modeled by the backbone (Eq 1), we extend the model to include further information of relevance to improve HAM detection, namely phylogenetic information and predictions of epitope peptide processing and MHC I affinity, as described in the following.

#### 2.2.1 Phylogeny

Viral strains have a common phylogenetic history. Thus substitutions are not independently and identically distributed, and therefore violate a common assumption of standard statistical methods. In fact, Bhattacharya et al. (2007) demonstrated the importance of correcting for the phylogenetic structure in identifying HLA associations.

A popular approach in phylogeny-aware regression of binary variables is to estimate an additional multivariate normally distributed intercept, where the covariance matrix is based on the branch lengths of a given phylogenetic tree (Ives and Garland, 2009, 2014). This approach turned out to be too computationally expensive in our model, hence we chose a strategy similar to the one in Carlson et al. (2008):

Consider a phylogenetic tree Ψ obtained from standard maximum likelihood methods for a given multiple sequence alignment. We are interested in estimating *P*(*y*_*ik*_ = 1|Ψ), that is, the probability of observing the substitution *k* in sequence *i* based on the underlying phylogenetic model. A quantity that can be readily computed using phylogenetic software like RAxML-NG (Kozlov et al., 2019) is *P*(Ψ|*y*_*ik*_ = 1). For this, we keep the tree topology fixed, annotate the tree with the binary observations *y*_*ik*_ at its leaves and optimize the branch lengths. *P*(Ψ|*y*_*ik*_ = 1) is then the likelihood of the annotated phylogenetic tree. Similarly, we can also compute *P*(Ψ|*y*_*ik*_ = 0) by flipping the annotation of sequence *i* from 1 to 0 (keeping all other observations). With *P*(Ψ|*y*_*ik*_ = 1) and *P*(Ψ|*y*_*ik*_ = 0) known and the relative frequencies of 0 an 1 as priors, we can estimate *P*(*y*_*ik*_ = 1|Ψ) by applying Bayes’ theorem. The estimated probabilities based on phylogeny are then included in the model as additional intercepts (second term of logistic argument):

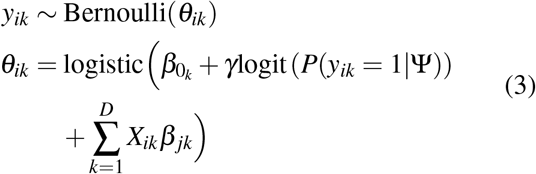

The logit transform is used because it cancels out with the logistic inverse link function. The phylogeny term acts as a baseline in absence of any HLA effects. As this baseline itself is not certain but subject to errors of the phylogenetic probabilities *P*(*y*_*ik*_ = 1|Ψ), we introduce an additional parameter *γ*.

#### 2.2.2 Inclusion of CTL epitope predictions

As outlined earlier, escape mutations often appear as HAMs. Given the underlying mechanism, it is not surprising that escape mutations are enriched in CTL epitopes, i.e. in those viral peptides presented by MHC I to TCRs (Bronke et al., 2013). This suggests that knowledge of epitope regions can be used to boost HAM detection. Fortunately, availability of large experimental datasets (Vita et al., 2019) has enabled the development of computational tools that predict with good accuracy the binding of peptides to MHC I molecules encoded by various HLA alleles (Mei et al., 2020).

Not only mutations in CTL epitopes can lead to failure to present epitopes to T cell receptors, but also mutations at epitope-flanking positions that interfere with pre-processing of peptides, notably proteasomal cleavage of viral proteins (Milicic et al., 2005; Gall et al., 2007).

In HAMdetector we use MHCflurry 2.0 (O’Donnell et al., 2020) to predict epitopes that are properly processed and presented by MHC I. For this, we create an input matrix of dimensions *R*×*D*, where *R* is the number of evaluated substitutions and *D* is the number of observed HLA alleles in the dataset. The elements of this matrix are binary encoded and contain a 1 if that position is predicted to be in an epitope, and 0 otherwise. Given an amino acid sequence, MHCflurry provides a list of possible epitopes (9-13 mers) and HLA allele pairs and calculates a rank based on comparisons with random pairs of epitopes and HLA alleles. For the binarization we use the rank threshold of 2% suggested by MHCflurry.

We use epitope prediction as information about the expected degree of sparsity, i.e. if we know that there is an epitope restricted by a given HLA allele at that location, we expect that this HLA allele is more likely to be associated with substitutions at that position than the other HLA alleles. This idea is implemented by increasing the scale of the local shrinkage parameters *λ*_*jk*_ depending on epitope information:

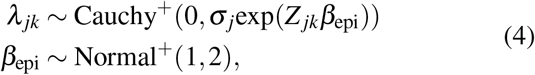

where *Z*_*jk*_ is 1 if HLA allele *j* is predicted to restrict the alignment position corresponding to substitution *k*, and 0 otherwise. The parameter *β*_epi_ governs the increase in scale of the corresponding local shrinkage parameters. The larger the estimated values of *β*_epi_ are, the more likely it is to see non-zero regression coefficients for these HLA alleles.

#### 2.2.3 Sparsity-inducing priors

Sparsity-promoting priors (Piironen and Vehtari, 2017b) can drastically improve predictive performance, because the model is better able to differentiate between signal and noise. These priors convey the a priori expectation that most coefficients in a regression model are close to 0, i.e. that non-zero coefficients are sparse. This assumption is likely correct for HAMs: the dominating mechanism that leads to HLA association of mutations is probably selection of mutations that mediate escape from MHC I presentation of epitopes; however, we know that these epitopes are sparse, i.e. the number of actual epitopes that are restricted by a given HLA allele is typically small compared to the number of all conceivable epitopes. Thus, for most pairs of HLA allele and substitution, the association is likely truly zero. Note that this reasoning does not preclude associations outside of epitopes as sometimes observed for compensatory mutations (Ruhl et al., 2011) but just implies that these are more rare.

There is a range of sparsity-promoting priors with slightly different properties. They share the common structure of placing most probability mass very close to 0, with heavy tails to accommodate the non-zero coefficients. For our model, we use the so-called regularized horseshoe prior (Piironen and Vehtari, 2017b), which is an improvement of the original horseshoe prior presented by Carvalho et al. (2010), in that it additionally allows some shrinkage for the non-zero coefficients. The original horseshoe prior is given by:

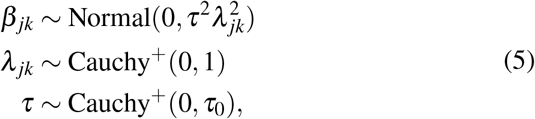

where *β*_*jk*_ are the regression coefficients; *τ* and *λ*_*jk*_ are the so-called global and local shrinkage parameters, respectively; Cauchy^+^ is the positively constrained Cauchy distribution; *τ*_0_ is the overall degree of sparsity. Shrinkage of the non-zero coefficients in the regularized horseshoe prior is achieved by replacing 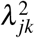 with 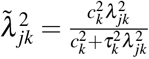, where the additional parameter *c* governs the magnitude of shrinkage for the non-zero coefficients.

With Eq 5 the global shrinkage parameter *τ* is typically very small and shrinks most of the regression coefficients close to 0, whereas the local shrinkage parameters *λ*_*jk*_ can occasionally be very large to allow some coefficients to escape that shrinkage.

The overall degree of sparsity *τ*_0_ can be chosen based on the expected number of non-zero coefficients (Piironen and Vehtari, 2017a).

### 2.3 Full model specification

The full specification of the HAMdetector model is:

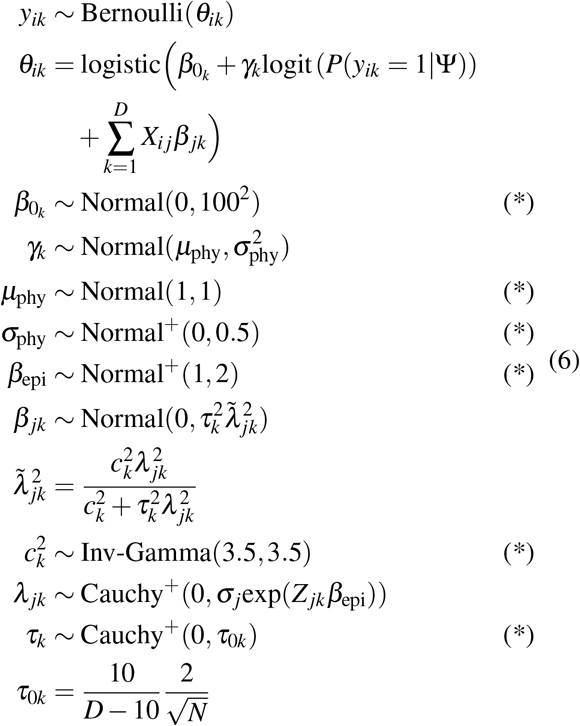

where *N* is the number of available annotated sequences.

The full model specification includes some aspects that were not covered in the previous sections. In particular, the overall phylogeny-weight *γ* in Eq 1 is replaced in the full model by hierarchically modelled *γ*_*k*_, which allows partial pooling across substitutions (even with a global parameter *γ* the model works reasonably well). The final additional parameters 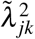 and *τ*_0*k*_ are explained in detail in Piironen and Vehtari (2017b). Briefly, 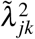 allows some regularization for the non-zero coefficients and the parameterization of *τ*_0*k*_ allows to place a prior on the expected number of non-zero coefficients. This is particularly useful for logistic regression models, as some shrinkage helps to deal with issues of separability and collinearity that commonly occur with logistic regression models.

#### 2.3.1 Prior justification

Prior distributions are labeled with an asterisk in Eq 6. They are weakly informative, which means that they effectively limit posteriors to realistic magnitudes of parameters. One exception to this are the intercepts 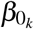, which are essentially flat because they are well identified by the data alone.

The hierarchical mean and standard deviation of the phylogeny coefficients *γ*_*k*_ place most probability mass on *γ*_*k*_ values around 1. In absence of any HLA effects, a *γ*_*k*_ = 1 would mean that the estimate for the probability of observing substitution *k* is identical to the probability based on the phylogenetic model. This treats phylogeny as a baseline, and any observations not attributed to phylogeny must be explained by HLA alleles or noise.

The prior on 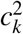 implies a Student-t prior with 7 degrees of freedom and a scale of 1 on the non-zero HLA regression coefficients *β*_*jk*_. A Student-t prior with these parameters is a reasonable default choice for logistic regression models (Piironen and Vehtari, 2017b).

The value of *τ*_0*k*_ implies 10 effective non-zero HLA regression coefficients per substitution. The rationale behind this parameterization is again outlined in Piironen and Vehtari (2017b). The value of 10 corresponds to a generously estimated magnitude based on available HIV epitope maps (Yusim et al., 2018). The model is also parameterized in a way that assumes an equal degree of sparsity across all alignment positions a priori. We also tried to model *τ*_*k*_ hierarchically, but observed sampling issues due to the resulting unfavorable geometry of the posterior.

### 2.4 Model implementation

A Julia (Bezanson et al., 2017) package is available at https://github.com/HAMdetector/Escape.jl to run the model on custom data. Due to restrictions of dependencies (MHCflurry and RAxML-ng), HAMdetector is currently only available on Linux, but can be run on Windows using the Windows Subsystem for Linux (WSL2). All models were implemented in Stan 2.23 (Stan Development Team, 2021), a probabilistic programming language and Hamiltonian Monte Carlo sampler for efficient numerical computation of posterior distributions. The Stan code is available in two versions: One optimized for readability and one optimized for speed by utilizing Stan’s multithreading and GPU capabilities.

### 2.5 Model diagnostics

#### 2.5.1 Convergence diagnostics

We use the split-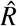 convergence diagnostic to identify Markov chain convergence issues (Gelman and Rubin, 1992; Gelman et al., 2013). We require a value of 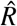 below 1.01 for all model parameters. Additionally, we require that the effective sample size N_eff_ (Stan Development Team, 2021) is above 500 for all model parameters and that sampling occurs without any divergent transitions (Betancourt, 2017).

#### 2.5.2 Posterior predictive checks

In posterior predictive checks, we simulate new data from the inferred posterior distribution and the likelihood, and we compare these simulated data with representative real data (Gabry et al., 2019). A good model should predict data that are consistent with real data. This general idea was employed in two ways to test our models.

For a first posterior predictive check we used *calibration plots* (Fig. 1): two binned quantities were plotted against each other, the observed relative frequencies of substitutions *f* (*y*_*ik*_ = 1), and the predicted probabilities *P*(*y*_*ik*_ = 1|model). In such a plot, a well-calibrated model should yield points following the diagonal. Technically, all observations were first sorted by increasing estimated probability *P*(*y*_*ik*_ = 1|model) and grouped into *n* bins. For each bin, the fraction of observations with *y*_*ik*_ = 1 (observed event percentage) was then plotted against the midpoint of each bin. The cutpoints of the bins are indicated by error bars.

**Figure 1.**
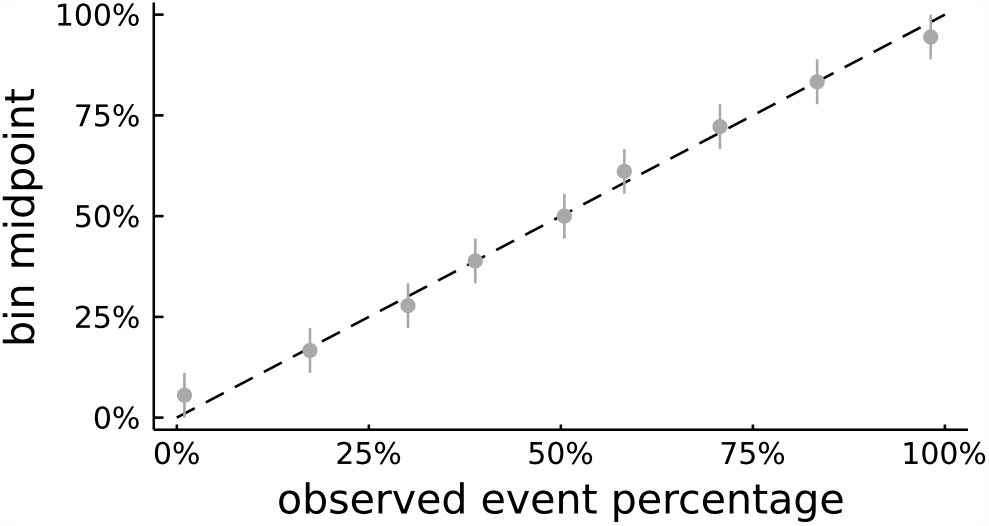
Calibration plot for the HBV PreC/core protein.

Second, we assessed the abilities of different models and methods to discover HAMs with *HAM enrichment plots*. These plots are based on the observation that CTL escape mutations are enriched in epitopes (Bronke et al., 2013). Hence, the degree by which methods for HAM prediction recover this trend is a measure of model performance. To implement this measure, we first ranked all evaluated substitutions according to their respective credibility of being a HAM, computed as integral of the marginal posterior *P*(*β*_*jk*_ *>* 0). For comparison with established methods, namely Fisher’s exact test and Phylogenetic Dependency Network (Carlson et al., 2008), ranked lists based on p-values were computed. Then we calculated for each rank *r* the accumulated number *N*_*e*_(*r*) of predictions of this rank or better ranks were located inside known epitopes. The higher the curve *N*_*e*_(*r*), the higher the enrichment of predicted HAMs in epitopes, see e.g. Fig. 2.

**Figure 2.**
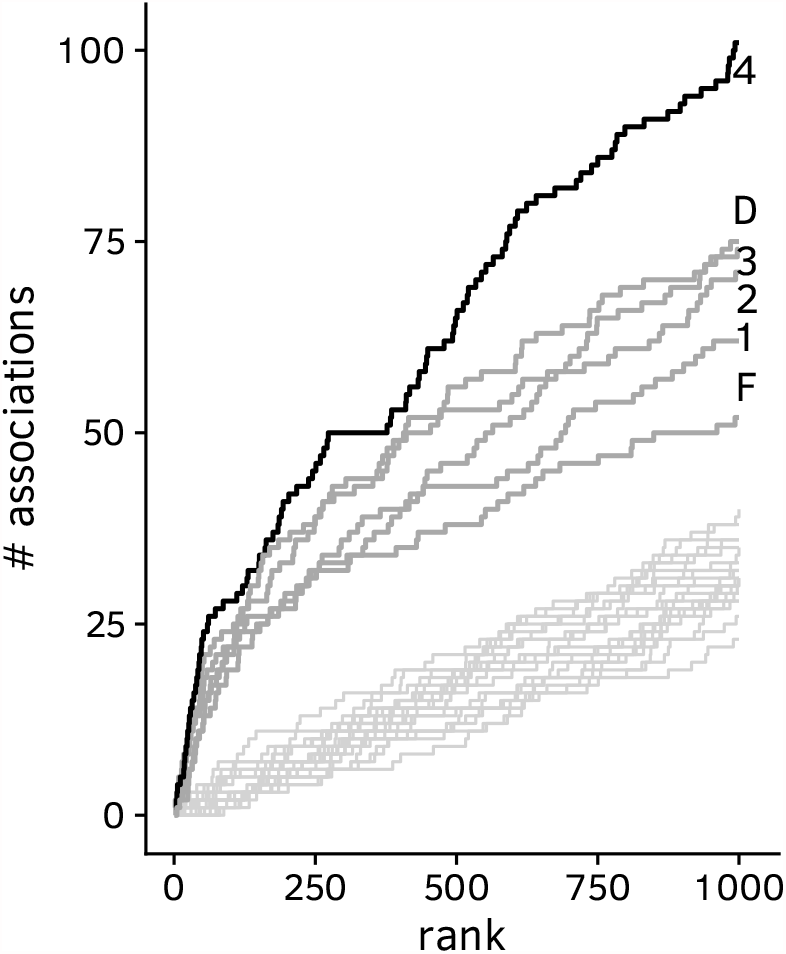
HAM enrichment plot for HBV preC/core protein: number *N*_*e*_ of associations inside the boundary of known epitopes vs. rank *r*. D: Phylogenetic Dependency Network; F: Fisher’s exact test; 1: simple logistic regression model with broad Student-t priors; 2: logistic regression model with horseshoe prior; 3: logistic regression model with horseshoe prior and phylogeny; 4: full model with epitope prediction. Unannotated gray lines at the bottom of the graph are HAM enrichment curves for random permutations of the list of HLA allele - substitution pairs and act as baselines.

#### 2.5.3 Leave-one-out cross-validation

Another performance measure is the ability to generalize to unseen data. To examine this ability for the different model variants we performed leave-one-out cross-validation (LOOCV), using the efficient Pareto-smoothed LOOCV Vehtari et al. (2016).

From the LOOCV, we obtain the Expected Log-Predictive Density (ELPD) 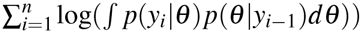 for samples *i* = 1, …, *n, i*th observation *y*_*i*_, data *y*_*i−*1_ with the *i*th data point left out, and model parameters *θ*. Thus, the ELPD is the average log predictive density of the observed data points based on the leave-one-out posterior distributions. This measure has the advantage over other performance measures like classification accuracy of not only taking into account the location of the predictive distribution (the number of correct predictions) but also the width, i.e. how confident the model is in its predictions.

### 2.6 Data

The model was fit with several datasets consisting of viral sequences paired to host HLA class I data:

- A large HIV dataset consisting of a subset of sequences from the HOMER (Brumme et al., 2007, 2008) cohort, the Western Australian HIV Cohort Study (WAHCS, Moore (2002); Bhattacharya et al. (2007)) and participants of the US AIDS Clinical Trials Group (ACTG) protocol 5142 (John et al., 2008) who also provided Human DNA under ACTG protocol 5128 (Haas et al., 2003) (total *N* = 1383). These data were in part also used in the Phylogenetic Dependency Network study (Carlson et al., 2008). The dataset contains sequences spanning the *gag, pol, env, nef, vif, vpr, vpu, tat* and *rev* genes.
- A set of 351 HIV sequences mostly spanning the *pol* gene from the Arevir database (Roomp et al., 2006).
- A set of 544 Hepatitis-B-Virus sequences (Timm and Walker, 2021) The dataset contains sequences of the preC/core, LHBs, Pol and HBx proteins.
- A set of 104 Hepatitis-D-Virus sequences containing the HDV-antigen (Karimzadeh et al., 2018).
- A set of 41 HIV sequences spanning the *gag* and *pol* genes.

Lists of known epitopes were gathered from the Immune Epitope Database (IEDB, Vita et al. (2019)).

### 2.7 Data preparation

For all sequences, we applied the following preparation steps:

1. For each dataset, the sequences were split into subsequences, either by protein or gene.
2. If not already present in this format, sequences were translated into their amino acid representations.
3. RAxML-NG (Kozlov et al., 2019) version 1.0.0 was used to generate a maximum likelihood phylogenetic tree for each gene/protein using the -model GTR+G+I option with all other parameters set to default values. If available, we used RNA or DNA sequences for this step, rather than protein sequences.

### 2.8 Data availability

The data underlying this article were provided by permission. Data will be shared on request to the corresponding author with permission of the respective co-authors.

## 3. Results

In order to understand what the different building blocks of HAMdetector contribute, we applied four different Bayesian models of increasing complexity to each dataset, starting with the standard logistic regression model (Eq 1), and adding then the further components, i.e. the horseshoe prior (Eq 5), phylogeny (Eq 3), and epitope prediction, resulting in the full model (Eq 6). For comparisons to existing methods, we also applied Fisher’s exact test and the Phylogenetic Dependency Network Carlson et al. (2008) to the same data.

### 3.1 Run times and convergence

For a standard office computer, run times of HAMdetector on the smaller HDV dataset were of the order of minutes and on the order of hours for the Hepatitis B dataset. For the large HIV dataset, the models were run overnight. Run times scale approximately linearly with the product *NK*, where *N* is the number of sequences and *K* is the number of substitutions. All model fits showed no signs of inference issues. In total, samples were drawn from four Hamiltonian Markov chains with 1000 iterations each after 300 warm-up iterations. The effective sample size exceeded 500 for all model parameters, 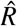 convergence diagnostic values were below 1.01 in all cases.

### 3.2 Posterior predictive checks

The model yields well-calibrated posterior predictive probabilities of substitutions. This is exemplified in Figure 1 for HBV core protein, but also holds true for the other datasets (Supplementary Figures “Calibration plots”).

The predictions of the tested models are enriched in epitopes over baselines for almost all tested datasets (Fig. 2 for HBV preC/core protein and Supplementary Figures “HAM enrichment plots” for other datasets). Although the relative and absolute performance varies by protein (see supplementary figure “HAM enrichment summary”), HAMdetector consistently outperforms all other methods in all but two datasets, and performs on-par with the other methods in these two cases. For the best ranked HAMs, Fisher’s exact test performs about as well as the HAMdetector backbone logistic regression model (model 1 in Fig. 2). Each of the following three model stages of HAMdetector increases HAM enrichment further. The horseshoe prior alone (model 2) is a drastic improvement over model 1, even though it does not include any specific external information. The logistic regression model with horseshoe prior works roughly as well as the Phylogenetic Dependency Network Carl-son et al. (2008), which includes much more information. Model 3 with its additional inclusion of phylogeny has higher enrichment than model 2, and finally, the full model 4 with the inclusion of epitope prediction leads to a further improvement. Note that model 4 only uses epitope *prediction* software and does not use any information of experimentally confirmed epitopes. The latter are here only used for model evaluation.

The Bayesian approach lends itself to incorporation of prior knowledge which usually helps in accurate modeling and prediction. In fact, a considerable effect is confirmed by the HAM enrichment plots with their ladder of improvements with increasing inclusion of information. It may be particularly surprising that the sparsifying horseshoe prior has such an impact although it does not use specific prior information. However, this is in principle the same mechanism as for the other information components: it is known that HAMs are sparse per HLA allele, and therefore supplying this information to the inference improves predictions. Figure 3 illustrates the effect of the sparsifying prior with an example, the substitution 11D in HIV integrase (Arevir dataset). There is no evidence for an association of HLA-A* 01 with this substitution, whereas for HLA-B* 44 the data is consistent with a strong association. The horseshoe prior has the effect of shrinking towards 0 specifically those regression coefficients with weak evidence of an association (A* 01 in Fig. 3). This reduces the standard error for the remaining coefficients, leading in our example to narrowed histogram for the association with B* 44 in the model with horseshoe prior.

**Figure 3.**
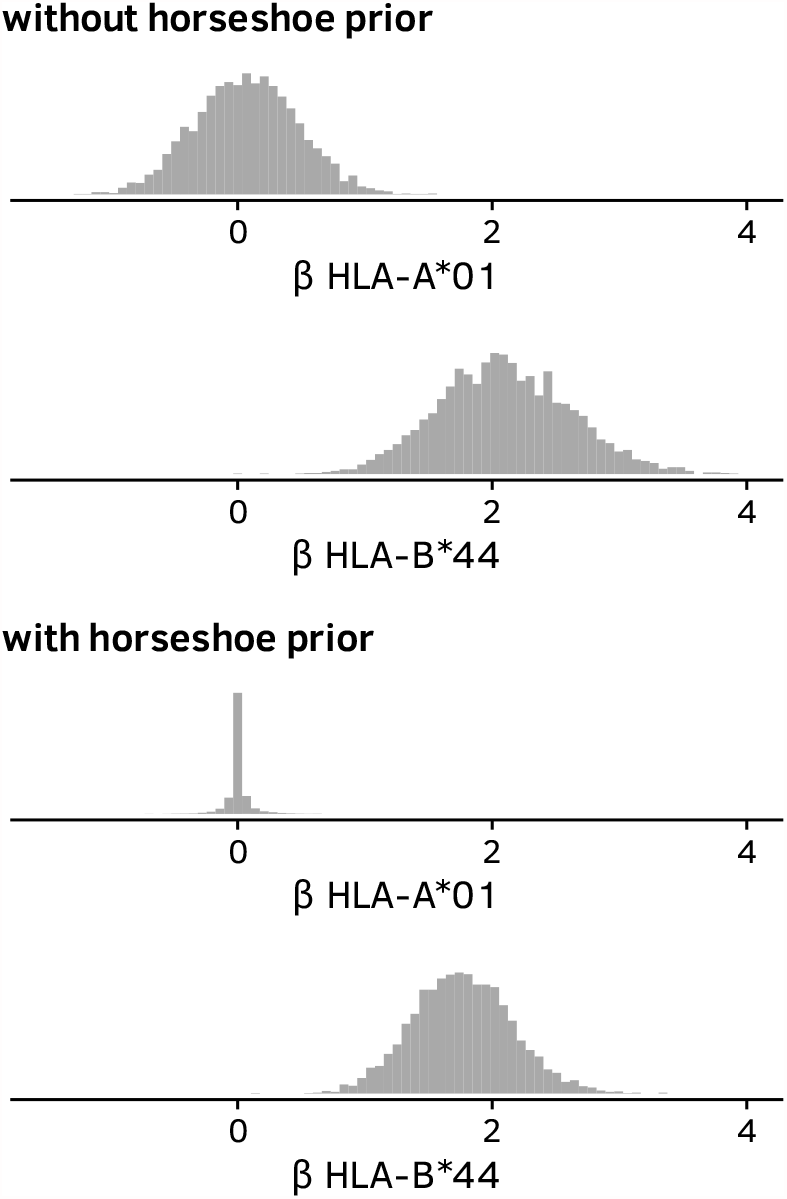
Marginal posterior distributions of regression coefficients for the association of substitution 11D of the HIV integrase with HLA alleles A* 01 and B* 44. Top half: inferred with logistic regression model, bottom half: inferred with logistic regression with sparsifying horseshoe prior.

### 3.3 Leave-one-out cross-validation

To quantify the ability of the four different model stages of HAMdetector to generalize to unseen cases, we computed the ELPD with Pareto-smoothed leave-one-out cross-validation. Table 1 shows results for the HBV preC/core protein in terms of ELPD changes with each new model stage. Each new model stage adds ELPD, i.e. is better at generalizing than the simpler model stages.

**Table 1.** ELPD changes as HAMdetector components are added. Data computed for HBV preC/core protein. All differences in ELPD are larger than several times the estimated standard error (column se_diff_), indicating that models that include more information have better predictive performance.

|  | ELPD<sub>diff</sub> | se<sub>diff</sub> |
| --- | --- | --- |
| logistic regression (baseline) | 0.0 | 0.0 |
| + horseshoe prior | 949.8 | 65.2 |
| + phylogeny | 4440.9 | 94.4 |
| + epitope prediction | 63.1 | 18.9 |

The model with horseshoe prior alone already has a much higher ELPD than the standard logistic regression model, even though it does not use any specific external data. This is because including the sparsity assumption allows the model to better separate signal from noise and the uncertainty of the close-to-zero coefficients does not propagate into uncertainty of predictions.

Including phylogeny further improves model performance a lot, as the assumption of independent and identically distributed data is replaced with specific information from the shared phylogenetic history.

While addition of sparsity and phylogeny has an effect on all substitutions and samples, epitope prediction only influences those substitutions that are restricted by a given HLA allele and only those samples that are annotated with the allele. Therefore, inclusion of epitope prediction does not improve ELPD as much as inclusion of phylogeny and the sparsity assumption. However, inclusion of epitope prediction is highly useful for determining which HLA alleles are associated with a substitution, as shown in the previous section.

### 3.4 HAMs in HDV as test case

The Hepatitis D Virus (HDV) dataset (Karimzadeh et al., 2019) is an excellent test case: we have (1) a set of paired HDV sequences and patient HLA alleles, (2) HAM predictions by Fisher’s exact test as implemented in SeqFeatR (Budeus et al., 2016), and (3) an in-vitro assay to quantify the effect of the predicted HAMs on IFN*γ* release of CD8^+^ T cells (IFN-*γ* production assays, Karimzadeh et al. (2019)). This allows us to see whether HAMdetector decreases the false positive rate in comparison to the simpler Fisher’s exact test, and we can make *bona fide* predictions on previously undetected HAMs. We have 15 HAMs predicted in HDV by Fisher’s exact test at significance level 5 *×* 10^*−*3^ (Table 2) as published (Karimzadeh et al., 2019). The corresponding p-values have no clear relation to experimental confirmation, i.e. p-values for confirmed HAMs are not generally lower than those of non-confirmed ones.

**Table 2.** List of HAMs predicted by Fisher’s exact test (FET). In the last column “+” and “-” mark experimentally confirmed or rejected HAMs, respectively; “?” below the horizontal line indicate untested bona fide predictions. “post.prob.” are posterior probabilities for positive associations computed with HAMdetector.

| substitution | allele | p-value (FET) | post. prob. | confirmed |
| --- | --- | --- | --- | --- |
| S170N | B*15 | $3 \cdot 10^{-8}$ | 0.99 | + |
| D101E | B*37 | 0.0002 | 0.96 | + |
| R105K | B*27 | 0.0011 | 0.93 | + |
| R139K | B*41 | 0.0034 | 0.92 | + |
| E47D | B*18 | 0.0027 | 0.90 | + |
| D33E | B*13 | 0.0001 | 0.86 | - |
| T134A | A*68 | 0.0045 | 0.82 | - |
| K43R | B*13 | 0.0021 | 0.77 | - |
| P89T | B*37 | 0.0011 | 0.75 | + |
| D47E | A*30 | 0.0010 | 0.76 | - |
| K113R | B*13 | 0.0043 | 0.76 | - |
| A107T | B*14 | 0.0028 | 0.70 | - |
| P49L | A*30 | 0.0031 | 0.63 | - |
| Q100L | B*13 | 0.0018 | 0.60 | - |
| D96E | B*13 | 0.0035 | 0.51 | - |
| E46D | A*02 | 0.0054 | 0.97 | ? |
| V81I | A*68 | 0.0073 | 0.97 | ? |
| K113N | B*08 | 0.0063 | 0.96 | ? |
| A71T | B*41 | 0.0065 | 0.96 | ? |
| L188I | A*68 | 0.0632 | 0.94 | ? |
| T95S | A*01 | 0.0285 | 0.93 | ? |
| D33E | A*03 | 0.0226 | 0.93 | ? |
| P49L | B*13 | 0.0035 | 0.92 | ? |
| A74S | A*68 | 0.0123 | 0.91 | ? |
| E29D | B*44 | 0.0559 | 0.91 | ? |
| D46E | B*57 | 0.0190 | 0.91 | ? |
| R88K | A*68 | 0.0123 | 0.91 | ? |
| T149P | B*52 | 0.0281 | 0.91 | ? |
| K43R | A*02 | 0.2158 | 0.90 | ? |
| N22S | B*08 | 0.0405 | 0.90 | ? |

For HAMdetector, we use in Table 2 the posterior probability of a positive regression coefficient (*P*(*β*_*jk*_ *>* 0) as measure for the confidence in having detected a HAM. HAMs with strong support have a posterior probability close to 1, associations with no support a probability close to 0.5 (corresponding to a regression coefficient centered around 0). The five predicted HAMs with top posterior probabilities (all ≥ 0.90) have all been experimentally confirmed. There is only one outlier with posterior probability 0.75 (P89T and B* 37).

HAMdetector strongly supports 15 substitution -allele pairs that have previously not been identified (question marks in last column of Table 2). All of them have association probabilities of 0.90 or higher, while their p-values from Fisher’s exact test exceed the significance level of 5 *×* 10^*−*3^ used in Karimzadeh et al. (2019). Given the superior performance of HAMdetector on the experimentally tested HAMs, these 15 bona fide predictions suggest that most true HAMs may still to be discovered. A striking example is K43R - A* 02 with a p-value of 0.22 in Fisher’s exact test but a HAM-probability of 0.90 and location inside an A* 02 restricted epitope.

### 3.5 Linkage disequilibrium

For three of the false positives proposed by Fisher’s exact test (Table 2), HAMdetector identifies associations with the same substitution but a different allele (P49L– B* 13 instead of P49L–A* 30; K43R–A* 02 instead of K43R–B* 13; and D33E–B* 13 instead of D33E–A* 03). One possible explanation for this observation is HLA linkage disequilibrium: If a certain HLA allele selects for a specific HAM and there is another HLA allele that co-occurs with that HLA allele, any method that relies on the statistical analysis of pairs of HLA allele and substitution alone will also detect these associations. Due to random sampling variation, the HLA allele that selects for a mutation might not necessarily have the strongest correlation. Inclusion of additional information like epitope prediction can help to identify associations that are otherwise confounded by noise.

Indeed, out of the 12 times P49L is observed in sequences annotated with A* 30, B* 13 is also present in 5 of those cases (Spearman’s rank correlation coefficient *ρ* = 0.5). A similar observation can be made for K43R and D33E, although the correlation between the respective alleles is much weaker. A* 30 and B* 13 have been shown to be in strong strong linkage disequilibrium (Brumme et al., 2007, supplementary table 2).

Figure 4 shows regression coefficients of the HLA alleles A* 30 and B* 13 for substitution P49L. With the simplest logistic regression model (model 1), both A* 30 and B* 13 have medium evidence of being associated with substitution P49L. However, with phylogeny and sparsity-promoting prior (model 3) both regression coefficients shrink close to 0 – the associations are not convincingly supported by the data. Using epitope prediction as additional source of information (model 4) allows to disentangle the association of the correlated alleles with P49L and identify B* 13 as likely associated with P49L. The association between P49L and A* 30 (predicted by Fisher’s exact test) remains shrunk towards 0.

**Figure 4.**
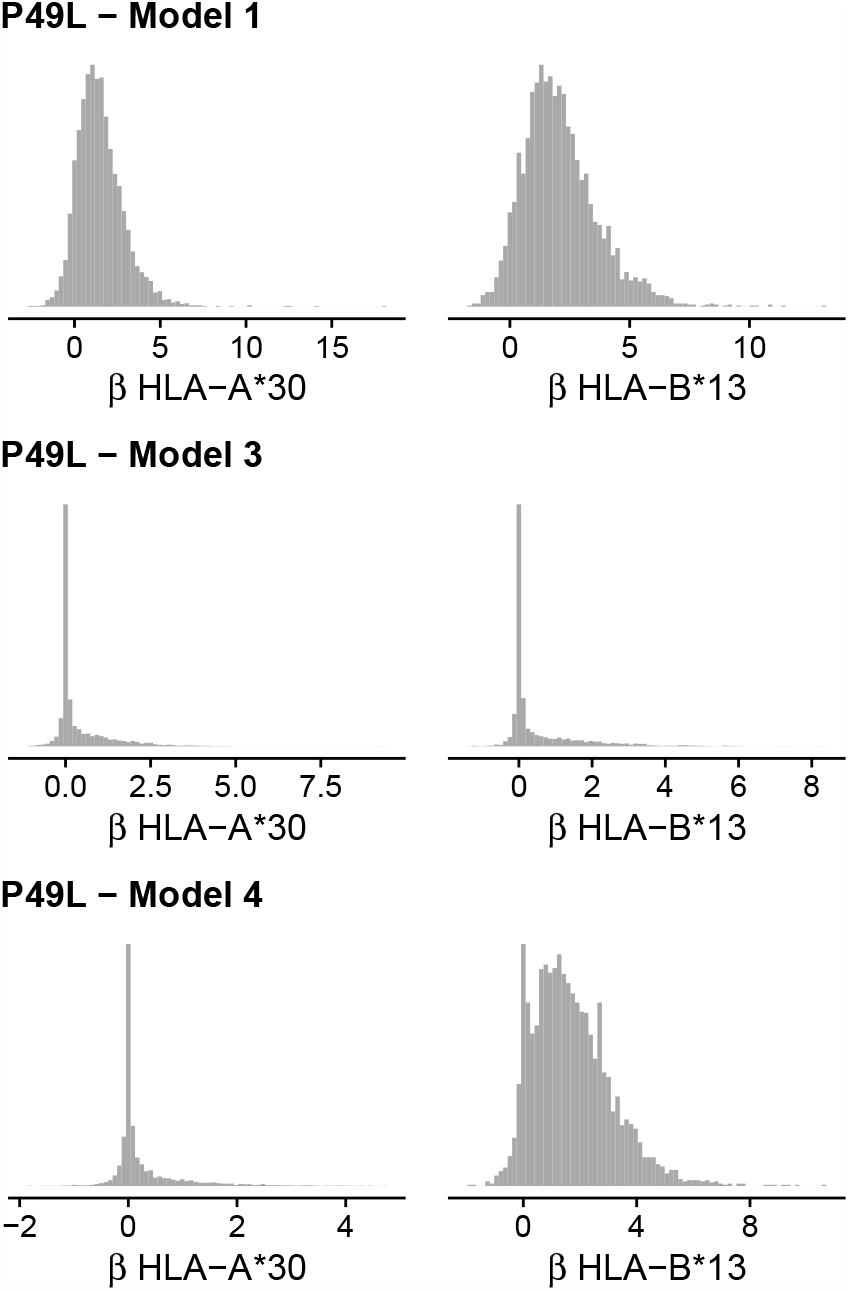
Marginal posteriors for the regression coefficients of A* 30 (left column) and B* 13 (right column) for substitution P49L with different model stages (rows).

### 3.6 HAMs outside epitopes

The epitope and processing predictions that HAMdetector uses are imperfect, as the underlying tools extrapolate binding affinities for new epitopes based on necessarily incomplete experimental data. Bayesian statistics provides a coherent framework to make use of imperfect data. In HAMdetector, this is achieved by an additional parameter *β*_epi_ that governs how strongly the model takes an apparent association between a substitution and the corresponding HLA allele into account. By default, the regression coefficients that quantify the strength of association between allele and substitution are shrunk towards 0, and only in the presence of considerable evidence in favor of an association (e.g. because the substitution often co-occurs with a certain HLA allele), this shrinkage is overcome by the observed data.

If the epitope prediction happens to be reliable, i.e. when the presence of a predicted epitope correlates strongly with the probability observing the substitution in a host with the respective HLA allele, the parameter *β*_epi_ is estimated to be large and less evidence by the sequence data is enough to escape the shrinkage and estimate a non-zero association between allele and substitution, compared to associations that do not lie inside a predicted epitope. Likewise, if the epitope prediction turns out to be non-reliable, *β*_epi_ is estimated to be close to 0 and the presence of a predicted epitope does not strongly affect the conclusions drawn from the sequence data.

However, it is important to consider that biologically relevant HAMs do not necessarily have to lie within or close to the boundary of an epitope. For instance, compensatory mutations can occur far away from the epitope they are associated with, as they might be the result of improved physical interactions with another amino acid in the folded, three-dimensional protein (Ruhl et al., 2011). Such compensatory mutations (Kelleher et al., 2001; Ruhl et al., 2011; Neumann-Haefelin et al., 2011; Schneidewind et al., 2008) can confer a strong selection advantage, e.g. by partially restoring replicative capacity that would otherwise be impaired by the exclusive presence of a certain HLA escape mutation.

We therefore also expect HAMs outside epitopes and one possible concern is that the model focuses too strongly on associations with substitutions that lie within the boundary of predicted epitopes.

Figure 5 shows posterior probabilities *P*(*β*_*jk*_ *>* 0) for substitution–HLA allele pairs as calculated by HAMdetector with (model 4) and without (model 3) epitope prediction. Each substitution–HLA allele pair is represented by a dot and colored according to whether or not that position lies within a predicted epitope. For substitutions that do not lie within a predicted epitope, both models provide similar estimates (points along the diagonal). However, some substitutions–HLA pairs that have only weak evidence of association in model 3 have strong support in model 4, which is explained by the additional evidence provided by epitope prediction. The figure shows that the model is still able to identify associations outside predicted epitopes and that epitope information augments evidence obtained from sequence data.

**Figure 5.**
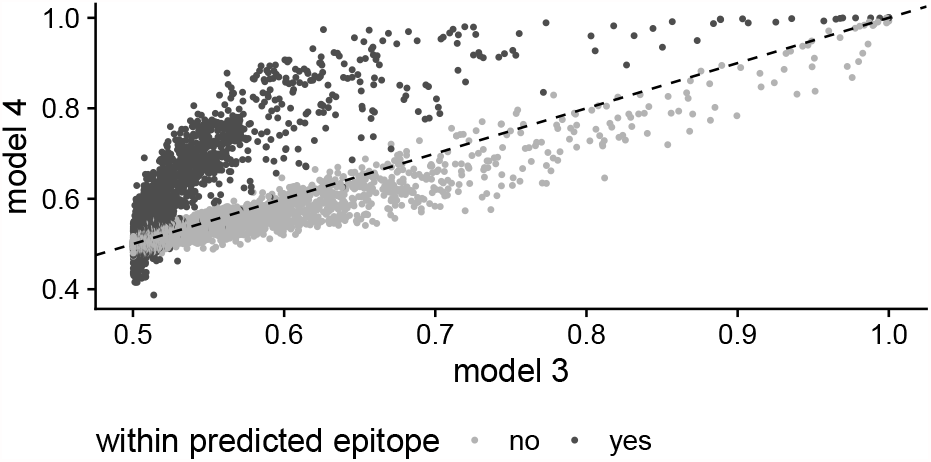
Integral of the marginal posterior *P*(*β*_*jk*_ *>* 0) for the HAMdetector model with epitope prediction (model 4) and without epitope prediction (model 3) for all substitutions in the preC/core protein (HBV dataset).

## 4. Discussion

HAMdetector follows a general paradigm of Bayesian modelling, namely to map all information that is available about a system of interest onto a probabilistic model, and then to apply Bayesian inference to learn about probable parameter values of that model, e.g. about *β*_*jk*_, the association of HLA *j* with substitution *k*. The more relevant information we infuse into the model, the sharper the inference. HAMdetector outperforms other methods as it includes an unprecedented amount of relevant information.

We have demonstrated that the logistic regression backbone is a platform that can be extended by model components that contribute new information. We have selected such modules guided by widely accepted knowledge, such as phylogeny or epitope location. However, even knowledge that is rarely stated explicitly may be helpful in inference, as in the case of sparsity of HLA associations. Since the included knowledge is generic for interactions of variable viruses with CTL immunity, HAMdetector performance does not depend on the virus.

Yet, HAMdetector is far from perfect. For instance, the outlier in Table 2 could point to missing information in HAMdetector. Another deficiency is that it currently works only with two-digit HLA alleles. We are currently exploring models for 4-digit HLA alleles that exploit partial pooling so that we can attenuate effects of the increased data fragmentation.

Another extension of our model would be to better account for phylogenetic uncertainty by using a Bayesian method to estimate a posterior distribution over possible tree topologies. The uncertainty over the tree topologies and the underlying parameters of the phylogenetic model would then propagate into uncertainty of the estimated probabilities *P*(*y*_*ik*_ = 1|Ψ). However, the good performance of the current version of HAMdetector makes it already a valuable tool for the study of interactions between viruses and T cell immunity.

## Funding

This work has been supported by Deutsche Forschungsgemeinschaft (grant HO 1582/10-1). ZLB is supported by the Canadian Institutes for Health Research (through project grant PJT-148621) and by the Michael Smith Foundation for Health Research (through a Scholar Award).

## Acknowledgements

We thank Drs. Mina John and Simon Mallal for providing data.

## Notes

https://github.com/HAMdetector/Escape.jl

### Competing Interest Statement

The authors have declared no competing interest.

