## Supplementary for "HAMdetector: A Bayesian regression model that integrates information to detect HLA-associated mutations"

Daniel Habermann[1,*], Hadi Kharimzadeh[2], Andreas Walker[3], Yang Li[4], Rongge Yang[4], Zabrina L. Brumme[5,6], Jörg Timm[3], Michael Roggendorf[7], and Daniel Hoffmann[1,8,9]

[1] *Bioinformatics and Computational Biophysics, Faculty of Biology, University of Duisburg-Essen, Essen, 45117, Germany*
[2] *Division of Clinical Pharmacology, University Hospital, LMU Munich, Munich, Germany*
[3] *Institute of Virology, Medical Faculty, University Hospital Düsseldorf, Heinrich-Heine-Universität, Düsseldorf, 40225, Germany*
[4] *AIDS and HIV Research Group, State Key Laboratory of Virology, Wuhan Institute of Virology, Chinese Academy of Science, Wuhan, P. R. China*
[5] *Faculty of Health Sciences, Simon Fraser University, Burnaby, Canada*
[6] *British Columbia Centre for Excellence in HIV/AIDS, Vancouver, Canada*
[7] *Insitute of Virology, School of Medicine, Technical University of Munich/Helmholtz Zentrum München, Munich, Germany*
[8] *Center of Medical Biotechnology, University of Duisburg-Essen, Essen, Germany*
[9] *Center for Computational Sciences and Simulation, University of Duisburg-Essen, Essen, Germany*

[*] To whom correspondence should be addressed.

### Calibration plots

#### HIV HOMER+

##### gag

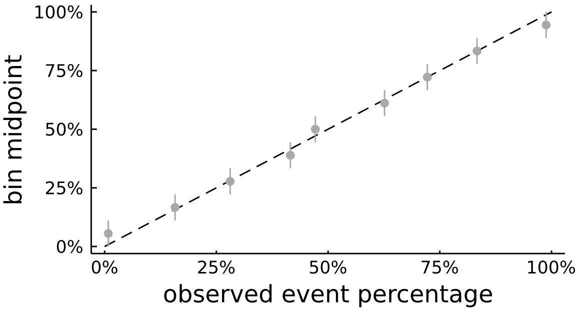


### gp41

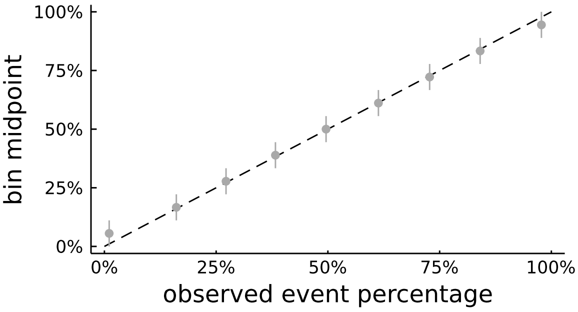


### gp120

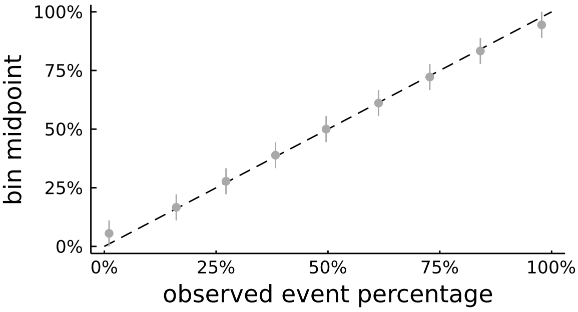


##### nef

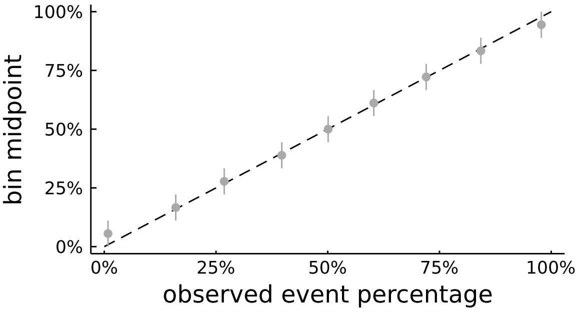


##### pol

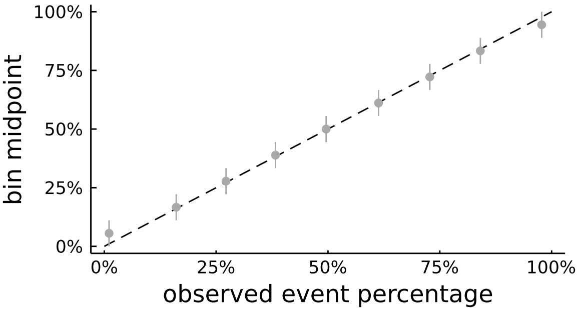


##### rev

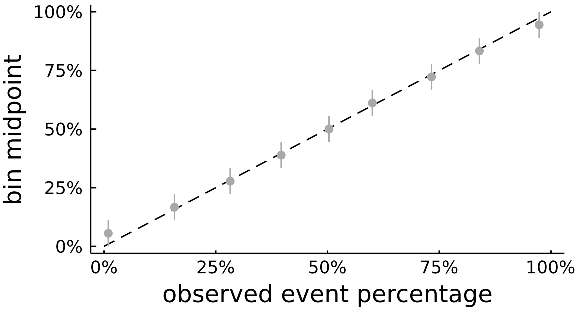


##### tat

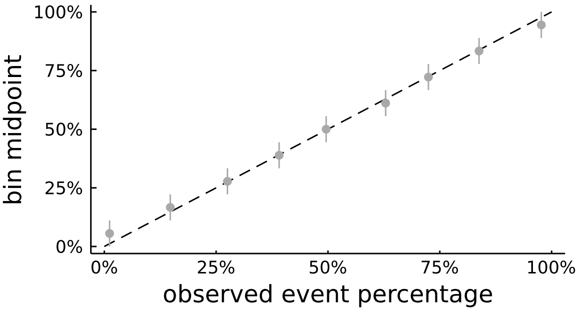


##### vif

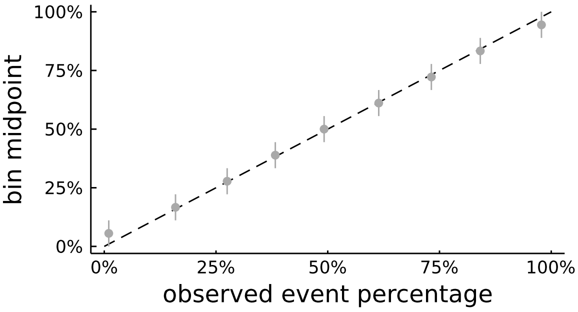

**vpr**

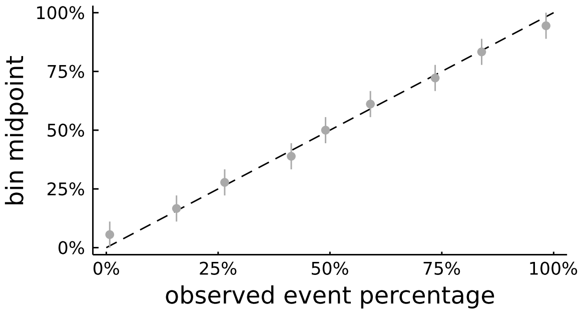
100%
75%
50%
25%
0%
bin midpoint
0%
25%
50%
75%
100%
observed event percentage

**vpu**

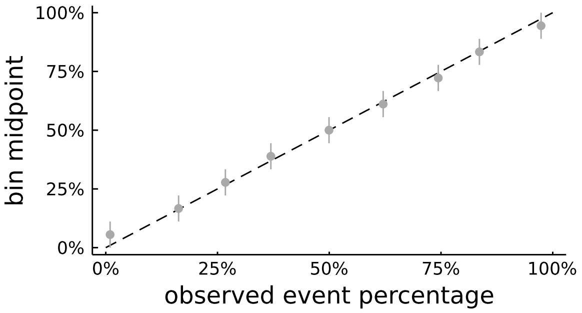
100%
75%
50%
25%
0%
bin midpoint
0%
25%
50%
75%
100%
observed event percentage

#### HBV

##### preC/core

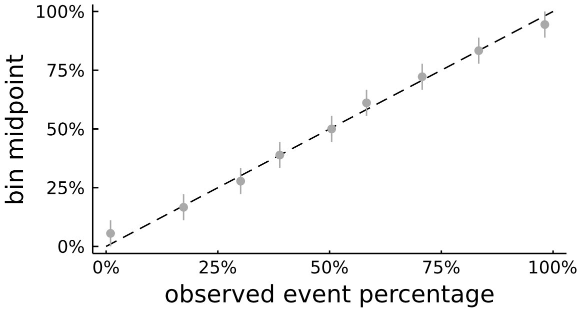


##### HBx

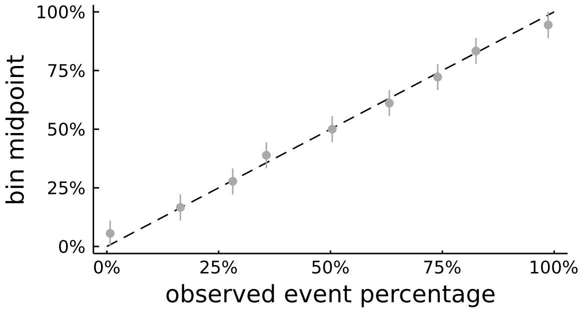


##### LHBs

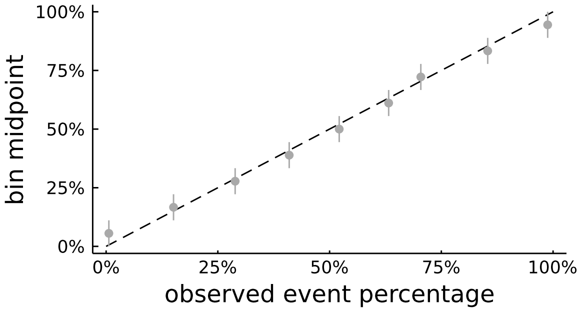


##### Pol

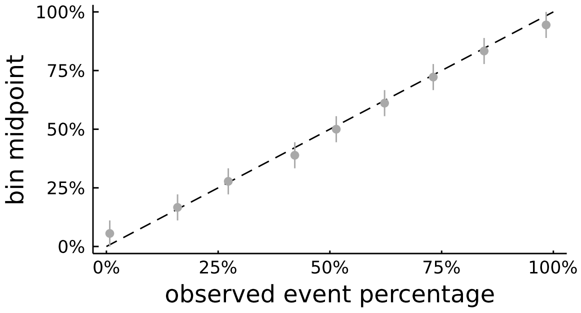


#### HDV

##### delta

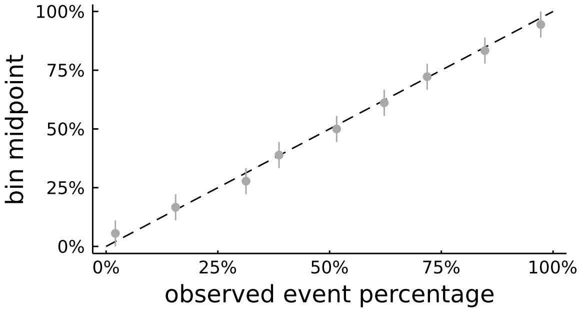

#### HIV Arevir

### gp120

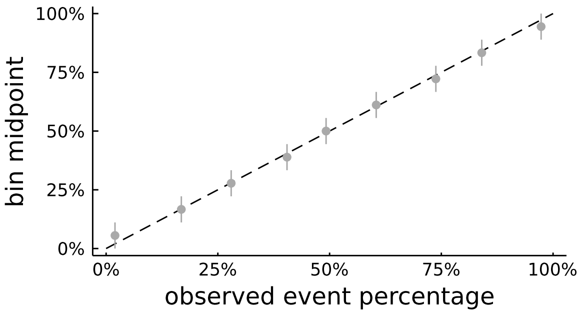


##### integrase

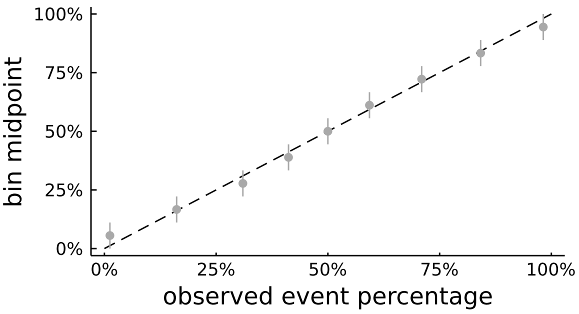


##### protease

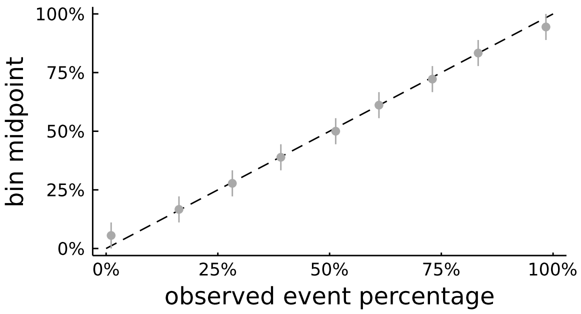


##### reverse transcriptase

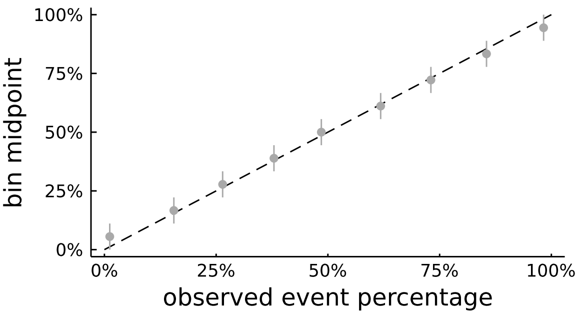

#### HIV

##### gag

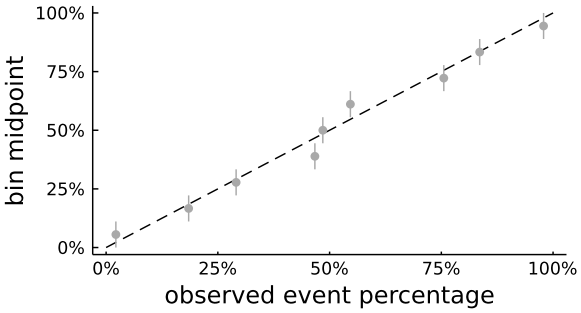


##### integrase

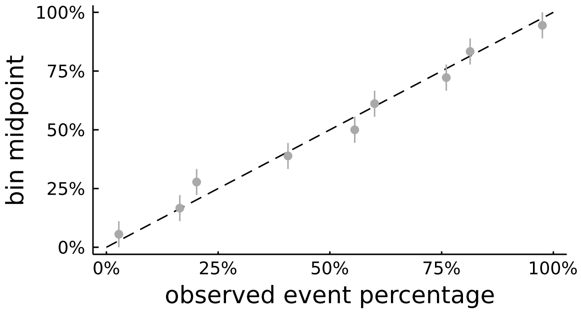


##### pol

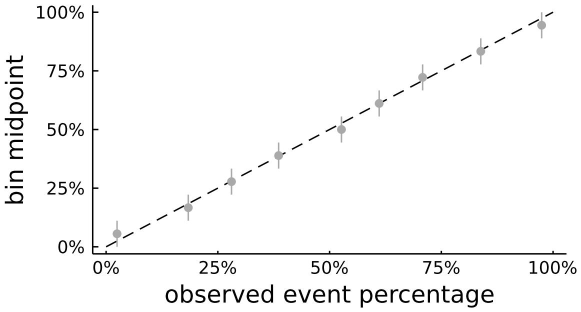


##### protease

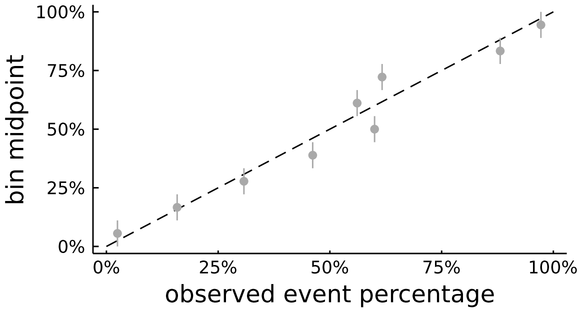


##### reverse transcriptase

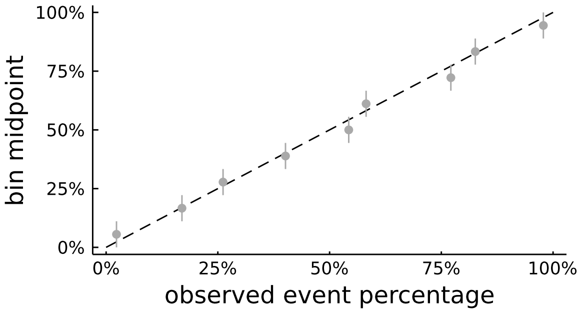

### HAM enrichment plots

#### HIV HOMER+

##### gag

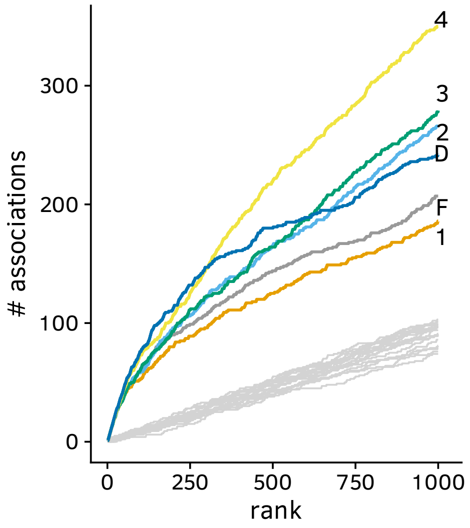


### gp41

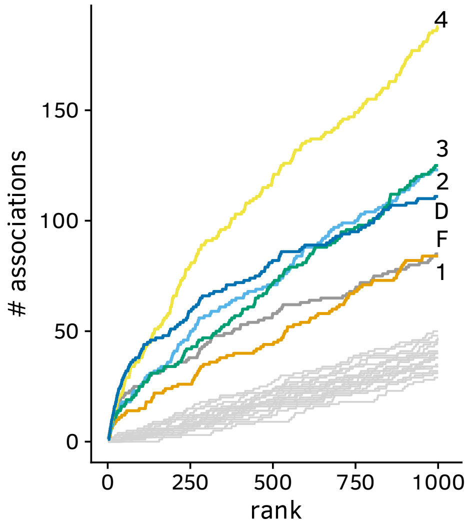


### gp120

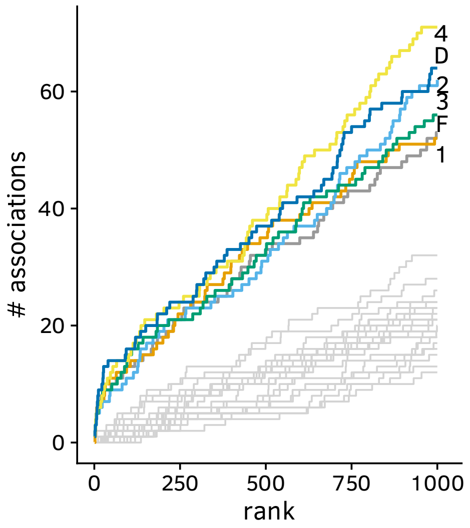


##### nef

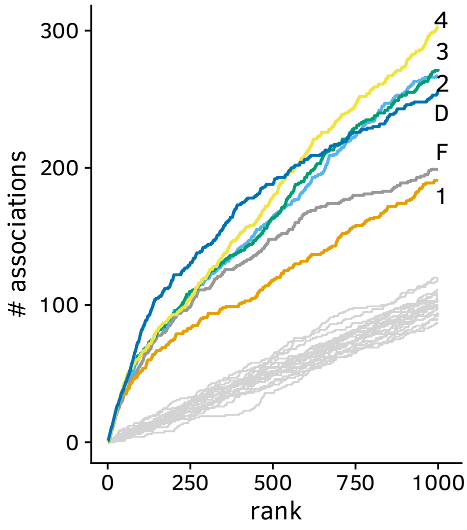


##### pol

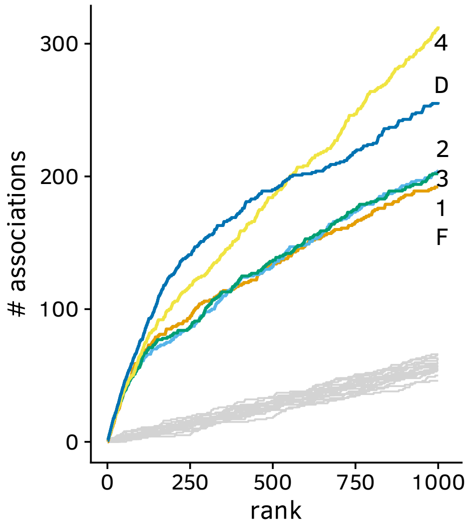


##### rev

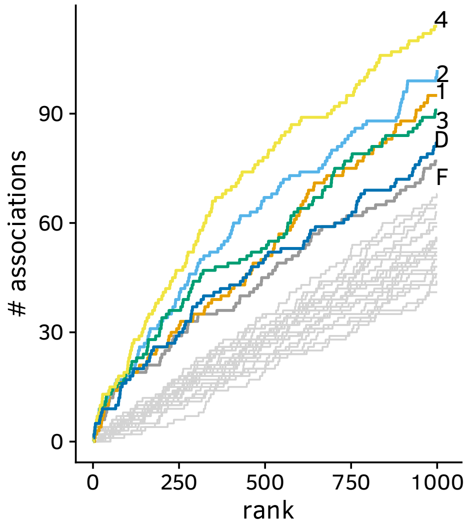

**tat**

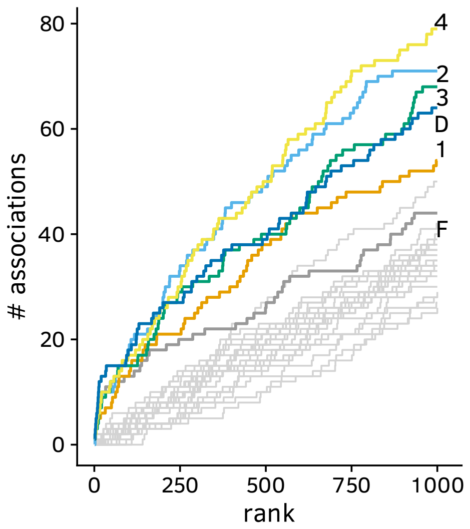


**vif**

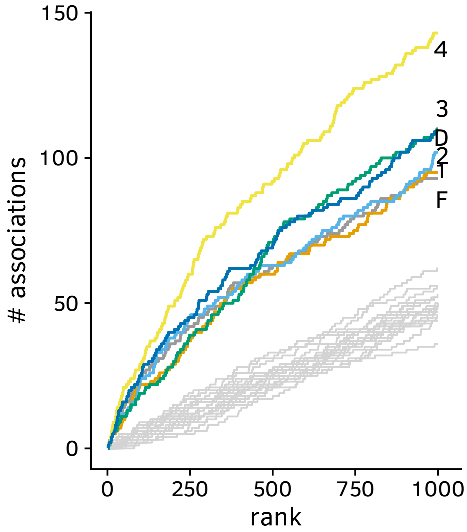


**vpr**

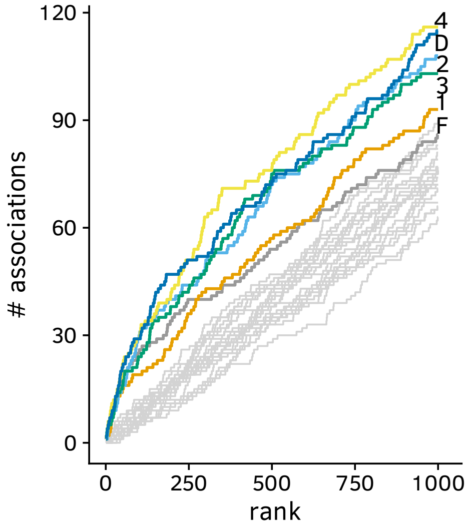


**vpu**

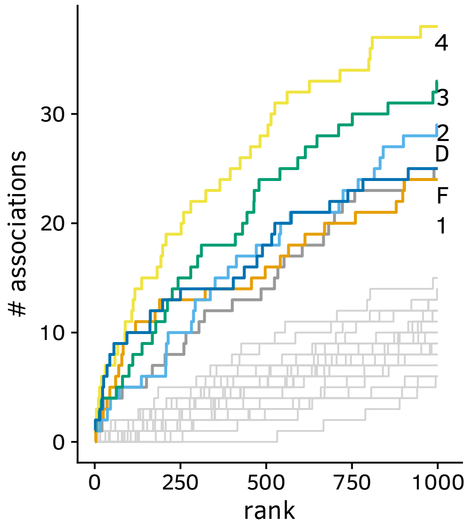

#### HBV

##### preC/core

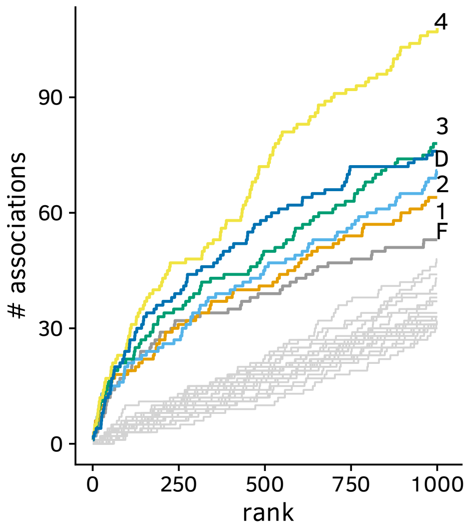


##### HBx

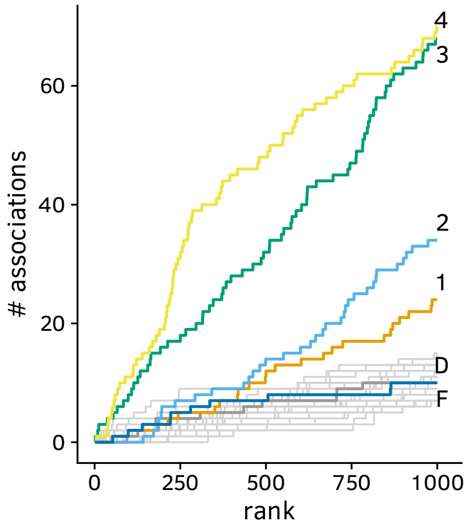


##### LHBs

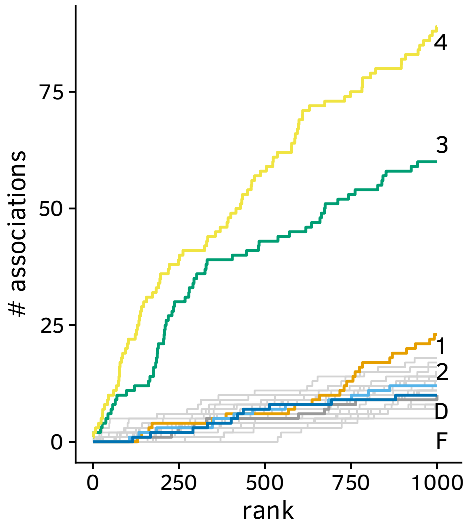


##### Pol

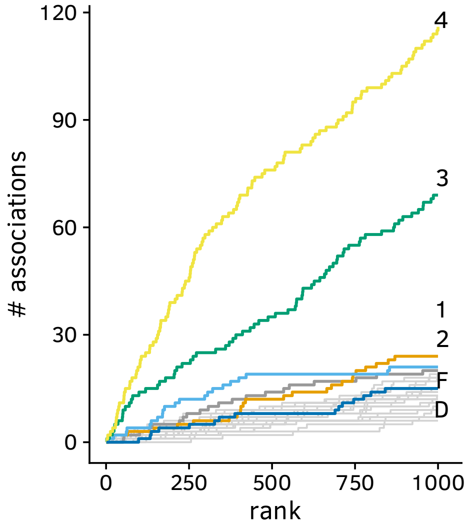

**HDV**

**delta**

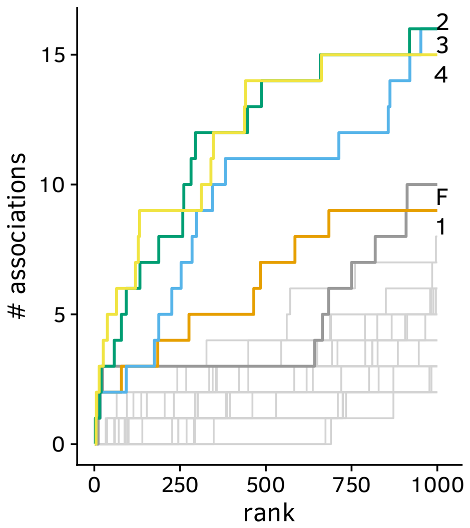

#### HIV Arevir

### gp120

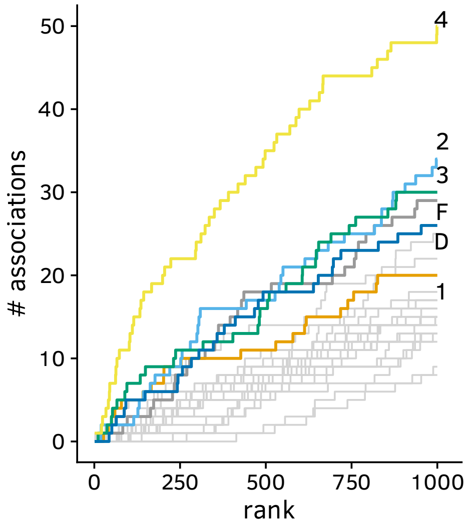


##### integrase

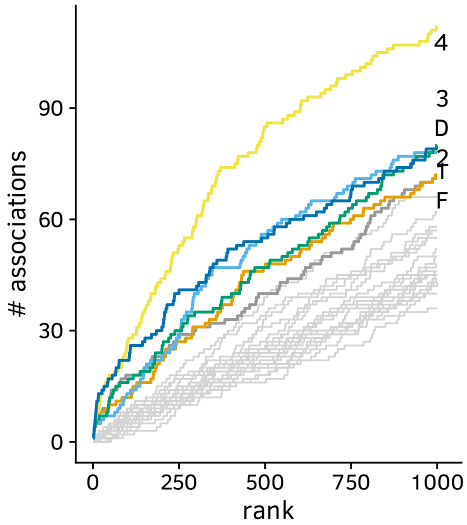


##### protease

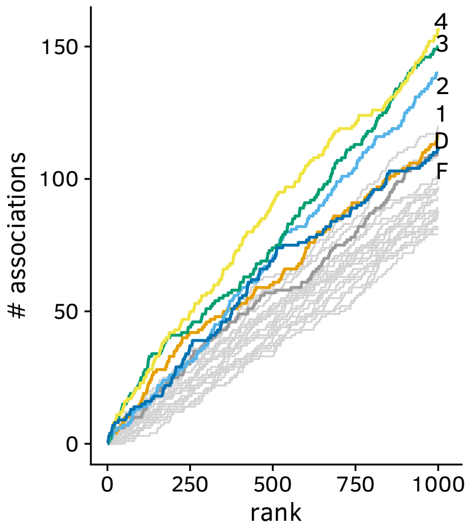


##### reverse transcriptase

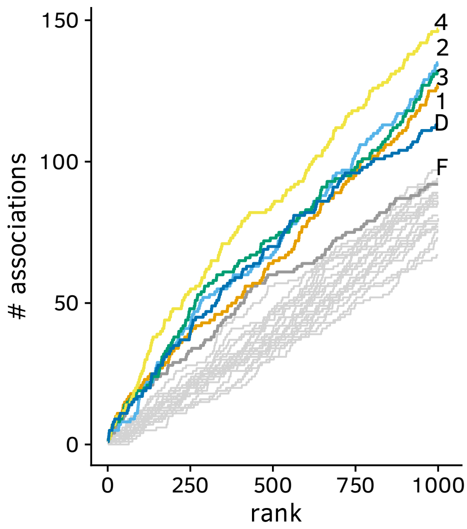

#### HIV CRF project

##### gag

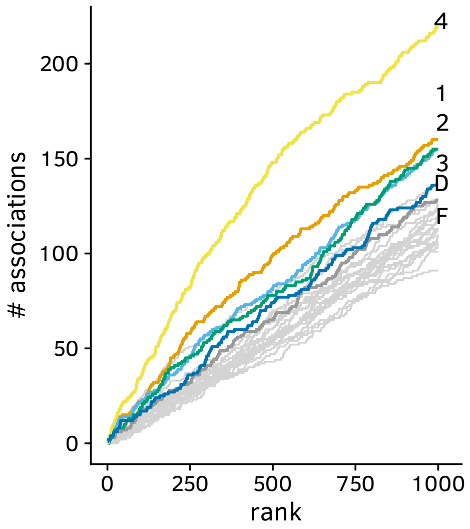


##### integrase

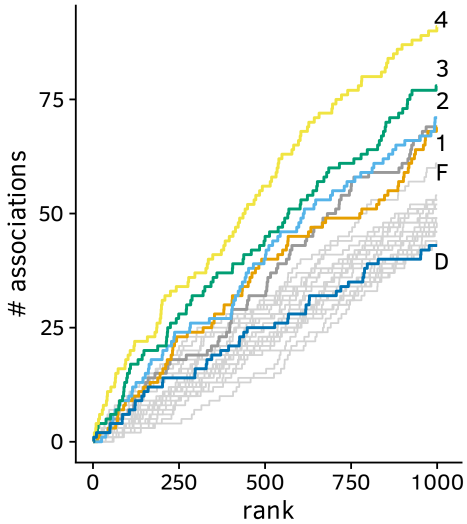


##### pol

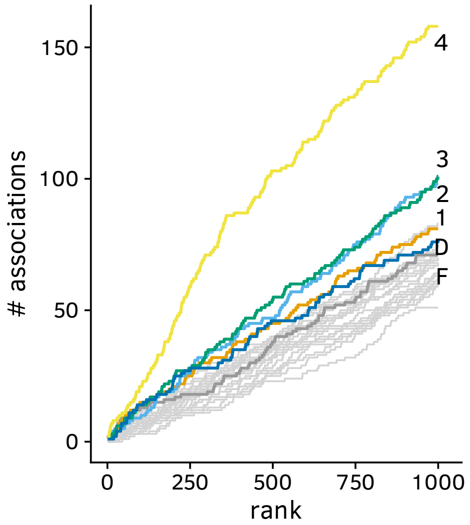


##### protease

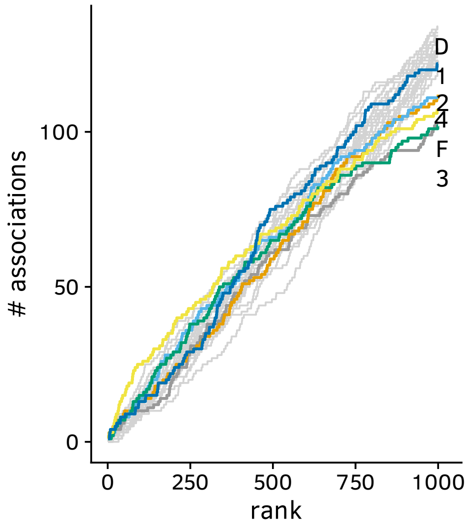


##### reverse transcriptase

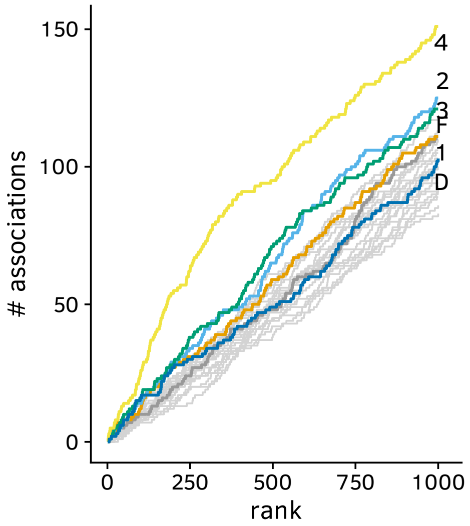

#### HAM enrichment summary

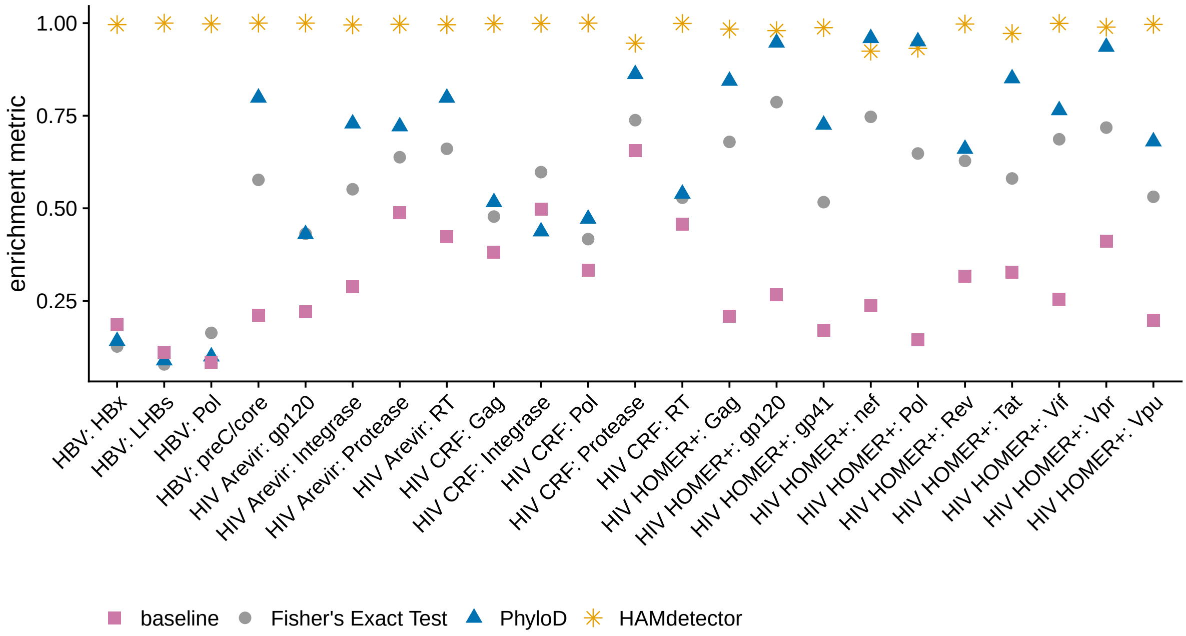


For each model and protein, an enrichment metric is computed according to the following algorithm: We first ranked all evaluated replacements according to heir respective credibility of being a HAM (integral of the marginal posterior $P(\beta_{jk} > 0)$ for HAMdetector, p-values for PhyloD and Fisher's exact testr). Let $r$ be the rank in that sorted list, e.g. rank 10 is the replacement–allele pair at position in that sorted list. Let $N_e(r)$ be the cumulative number of predictions of this rank or better that are located inside known epitopes. Let $m_r$ be the ratio between $N_e(r)$ for model $m$ and $N_e(r)$ for the top performing model. The enrichment metric $m$ is then computed as $m = \frac{1}{N}\sum_{r=1}^{N} m_r$ for ranks 1 to $N = 1000$.

### model 3 vs. model 4 scatter plots

#### HIV HOMER+

##### gag

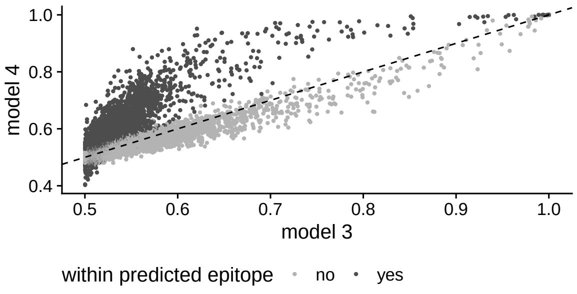


### gp41

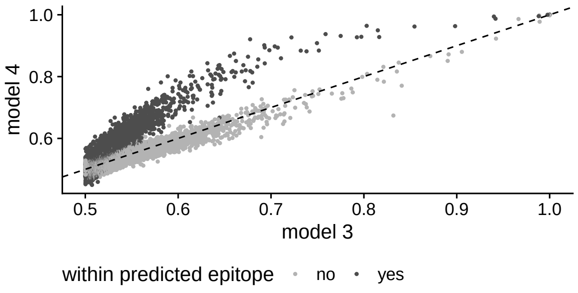


### gp120

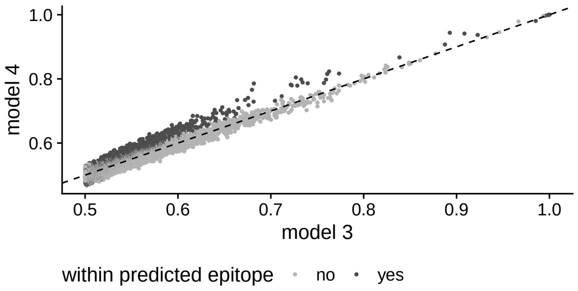


##### nef

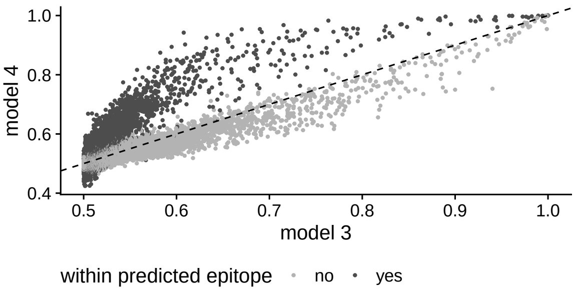


##### pol

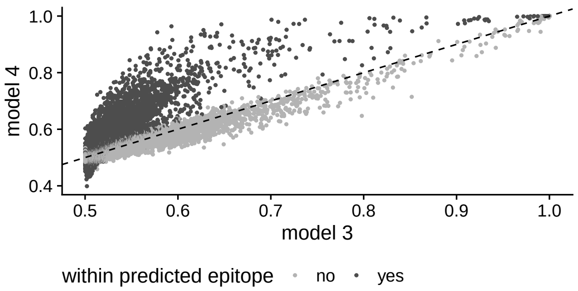


##### rev

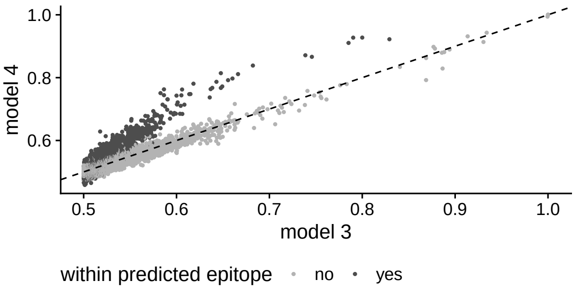

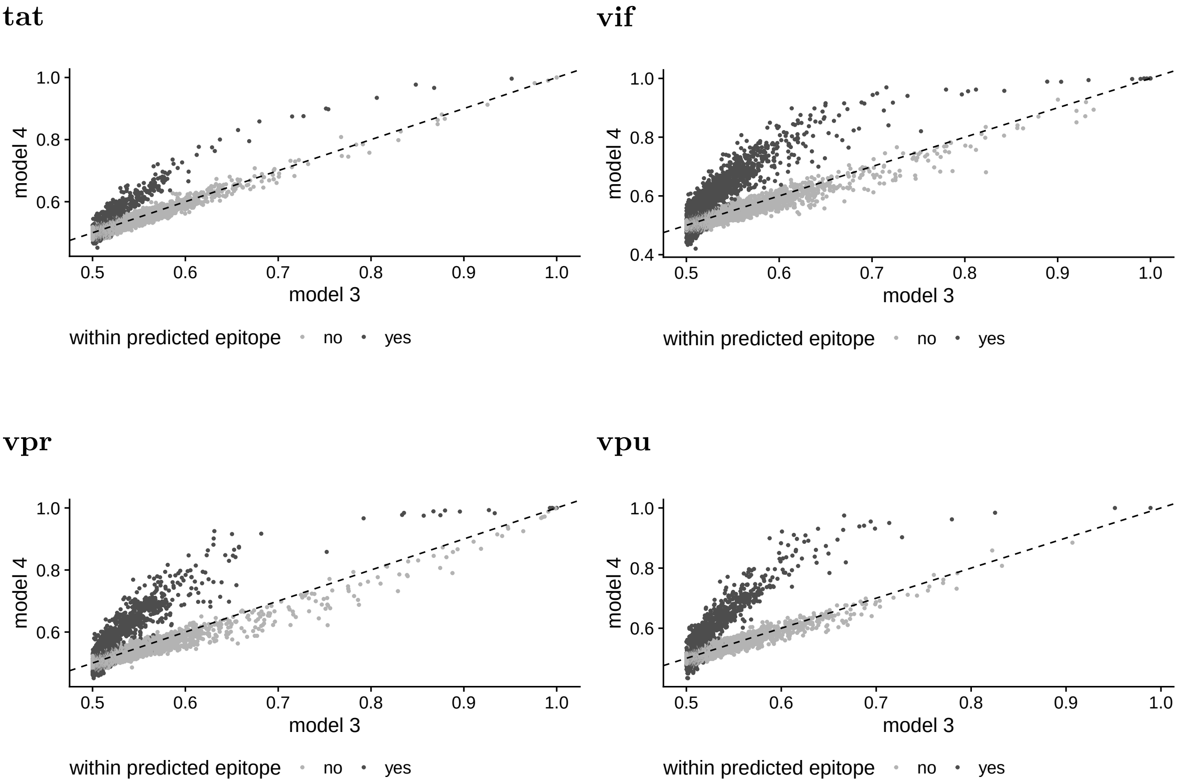
tat
vif
vpr
vpu
model 4
model 3
0.4
0.5
0.6
0.7
0.8
0.9
1.0
within predicted epitope
no
yes

### HBV

#### preC/core

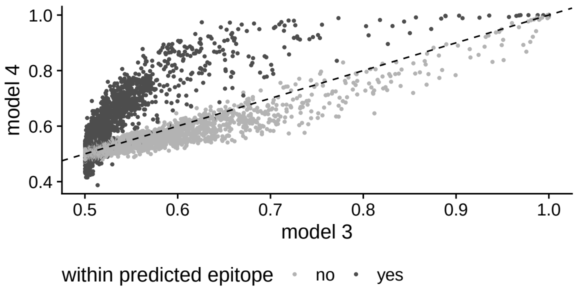


#### HBx

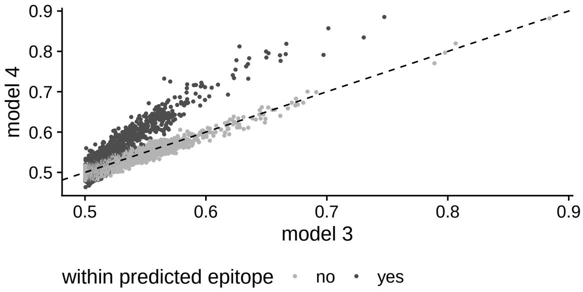


#### LHBs

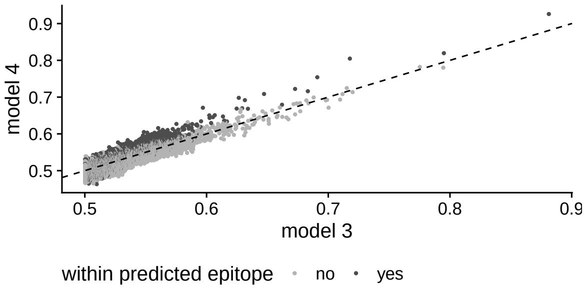


#### Pol

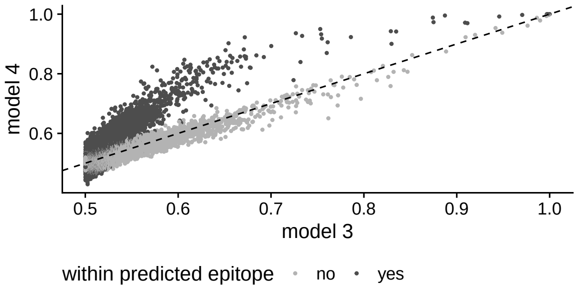


### HDV

#### delta

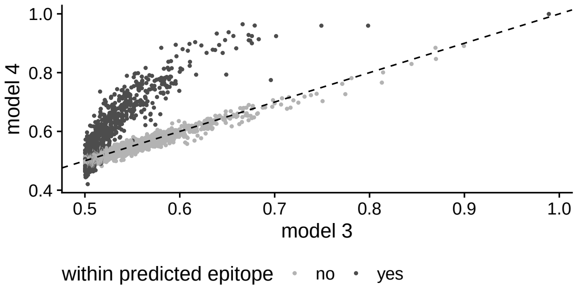

### HIV Arevir

## gp120

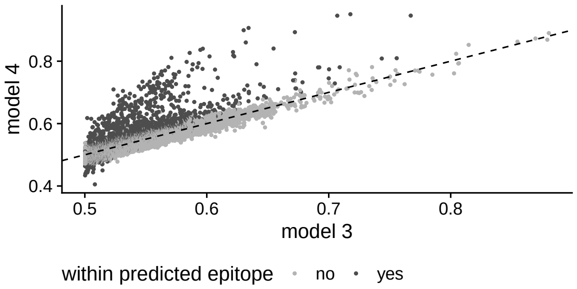


#### protease

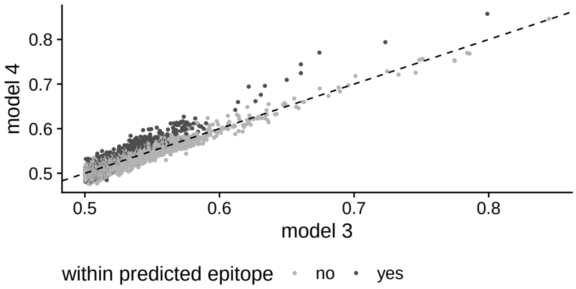


#### reverse transcriptase

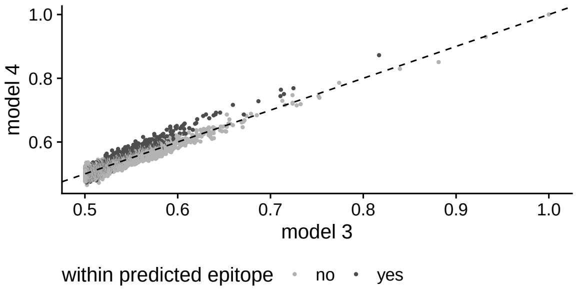

#### HIV CRF project

##### gag

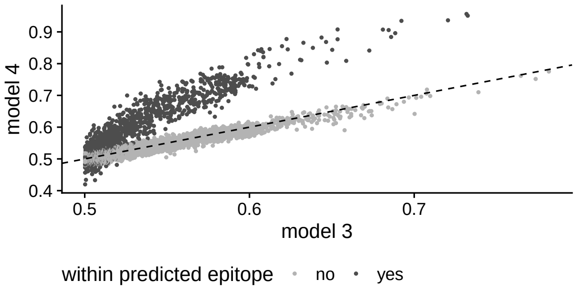


##### integrase

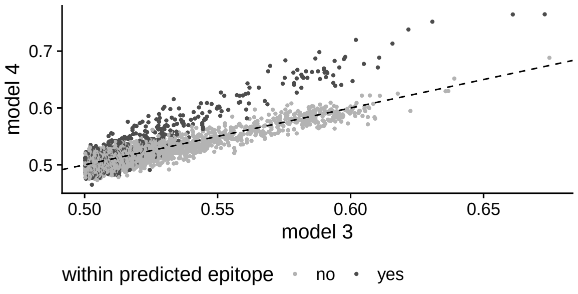


##### pol

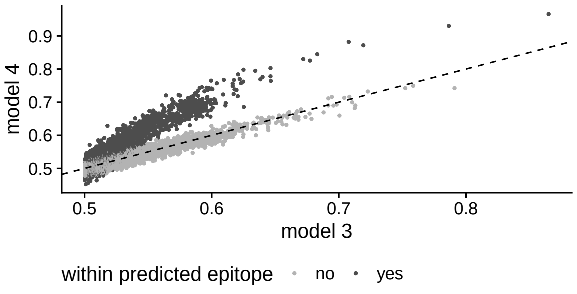


##### protease

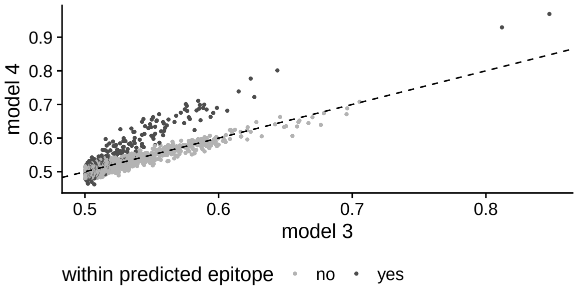


##### reverse transcriptase

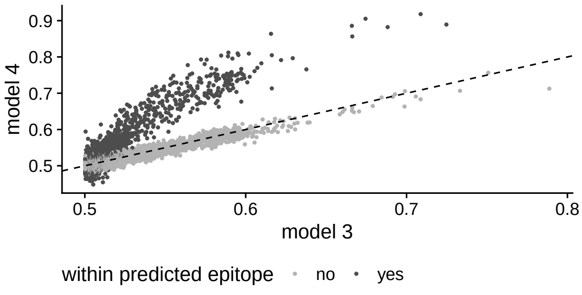

### LOO comparisons

#### HIV HOMER+

##### gag

| | $elpd_{diff}$ | $se_{diff}$ |
|---|---|---|
| logistic regression (baseline) | 0.0 | 0.0 |
| + horseshoe prior | 741.4 | 46.3 |
| + phylogeny | 3613.5 | 90.5 |
| + epitope prediction | 13.8 | 8.1 |

### gp41

| | $elpd_{diff}$ | $se_{diff}$ |
|---|---|---|
| logistic regression (baseline) | 0.0 | 0.0 |
| + horseshoe prior | 7626.7 | 140.4 |
| + phylogeny | 34068.3 | 276.5 |
| + epitope prediction | 64.7 | 16.3 |

### gp120

| | $elpd_{diff}$ | $se_{diff}$ |
|---|---|---|
| logistic regression (baseline) | 0.0 | 0.0 |
| + horseshoe prior | 14068.0 | 180.1 |
| + phylogeny | 13189.9 | 172.4 |
| + epitope prediction | -3.2 | 20.3 |

##### nef

| | $elpd_{diff}$ | $se_{diff}$ |
|---|---|---|
| logistic regression (baseline) | 0.0 | 0.0 |
| + horseshoe prior | 5065.2 | 119.4 |
| + phylogeny | 28433.4 | 255.1 |
| + epitope prediction | 51.5 | 20.1 |

##### pol

| | $elpd_{diff}$ | $se_{diff}$ |
|---|---|---|
| logistic regression (baseline) | 0.0 | 0.0 |
| + horseshoe prior | 9627.4 | 163.4 |
| + phylogeny | 40489.0 | 321.7 |
| + epitope prediction | 131.8 | 26.6 |

##### rev

| | $elpd_{diff}$ | $se_{diff}$ |
|---|---|---|
| logistic regression (baseline) | 0.0 | 0.0 |
| + horseshoe prior | 3400.4 | 97.6 |
| + phylogeny | 21225.3 | 215.9 |
| + epitope prediction | 17.0 | 9.9 |

##### tat

| | $elpd_{diff}$ | $se_{diff}$ |
|---|---|---|
| logistic regression (baseline) | 0.0 | 0.0 |
| + horseshoe prior | 3135.0 | 91.3 |
| + phylogeny | 19120 | 205.5 |
| + epitope prediction | 16.5 | 8.9 |

##### vif

| | $elpd_{diff}$ | $se_{diff}$ |
|---|---|---|
| logistic regression (baseline) | 0.0 | 0.0 |
| + horseshoe prior | 4343.0 | 108.1 |
| + phylogeny | 21254.6 | 221.3 |
| + epitope prediction | 38.8 | 16.2 |

**vpr**

| | $\text{elpd}_{\text{diff}}$ | $\text{se}_{\text{diff}}$ |
|---|---|---|
| logistic regression (baseline) | 0.0 | 0.0 |
| + horseshoe prior | 1714.2 | 71.1 |
| + phylogeny | 10488.2 | 155.3 |
| + epitope prediction | 23.4 | 12.4 |

**vpu**

| | $\text{elpd}_{\text{diff}}$ | $\text{se}_{\text{diff}}$ |
|---|---|---|
| logistic regression (baseline) | 0.0 | 0.0 |
| + horseshoe prior | 2877.5 | 85.4 |
| + phylogeny | 20004.2 | 199.7 |
| + epitope prediction | 50.1 | 12.8 |

#### HBV

##### preC/core

| | $elpd_{diff}$ | $se_{diff}$ |
|---|---|---|
| logistic regression (baseline) | 0.0 | 0.0 |
| + horseshoe prior | 971.5 | 65.2 |
| + phylogeny | 4427.8 | 94.6 |
| + epitope prediction | 65.7 | 19.5 |

##### HBx

| | $elpd_{diff}$ | $se_{diff}$ |
|---|---|---|
| logistic regression (baseline) | 0.0 | 0.0 |
| + horseshoe prior | 637.8 | 52.8 |
| + phylogeny | 8343.2 | 127.2 |
| + epitope prediction | 8.5 | 7.9 |

##### LHBs

| | $elpd_{diff}$ | $se_{diff}$ |
|---|---|---|
| logistic regression (baseline) | 0.0 | 0.0 |
| + horseshoe prior | 1744.2 | 84.6 |
| + phylogeny | 30120.1 | 223.8 |
| + epitope prediction | -7.6 | 14.7 |

##### Pol

| | $elpd_{diff}$ | $se_{diff}$ |
|---|---|---|
| logistic regression (baseline) | 0.0 | 0.0 |
| + horseshoe prior | 3157.7 | 127.7 |
| + phylogeny | 69561.1 | 343.5 |
| + epitope prediction | 98.8 | 19.0 |

#### HDV

##### delta

| | $elpd_{diff}$ | $se_{diff}$ |
|---|---|---|
| logistic regression (baseline) | 0.0 | 0.0 |
| + horseshoe prior | 455.5 | 37.5 |
| + phylogeny | 1746.9 | 63.1 |
| + epitope prediction | 32.1 | 12.2 |

#### HIV Arevir

### gp120

| | $elpd_{diff}$ | $se_{diff}$ |
|---|---|---|
| logistic regression (baseline) | 0.0 | 0.0 |
| + horseshoe prior | 5453.0 | 113.1 |
| + phylogeny | 12535.4 | 167.6 |
| + epitope prediction | -256.6 | 33.3 |

##### integrase

| | $elpd_{diff}$ | $se_{diff}$ |
|---|---|---|
| logistic regression (baseline) | 0.0 | 0.0 |
| + horseshoe prior | 1278.7 | 58.7 |
| + phylogeny | 6144.4 | 118.2 |
| + epitope prediction | 1.1 | 9.9 |

##### protease

| | $elpd_{diff}$ | $se_{diff}$ |
|---|---|---|
| logistic regression (baseline) | 0.0 | 0.0 |
| + horseshoe prior | 793.7 | 46.7 |
| + phylogeny | 3883.3 | 88.3 |
| + epitope prediction | -19.0 | 5.7 |

##### reverse transcriptase

| | $elpd_{diff}$ | $se_{diff}$ |
|---|---|---|
| logistic regression (baseline) | 0.0 | 0.0 |
| + horseshoe prior | 1653.7 | 65.7 |
| + phylogeny | 7034.2 | 127.5 |
| + epitope prediction | -6.1 | 10.7 |

#### HIV CRF project

##### gag

| | $elpd_{diff}$ | $se_{diff}$ |
|---|---|---|
| logistic regression (baseline) | 0.0 | 0.0 |
| + horseshoe prior | 352.2 | 36.4 |
| + phylogeny | 2893.1 | 73.1 |
| + epitope prediction | 1.5 | 13.6 |

##### integrase

| | $elpd_{diff}$ | $se_{diff}$ |
|---|---|---|
| logistic regression (baseline) | 0.0 | 0.0 |
| + horseshoe prior | 94.4 | 18.4 |
| + phylogeny | 564.8 | 36.7 |
| + epitope prediction | 0.0 | 6.5 |

##### pol

| | $elpd_{diff}$ | $se_{diff}$ |
|---|---|---|
| logistic regression (baseline) | 0.0 | 0.0 |
| + horseshoe prior | 543.9 | 42.4 |
| + phylogeny | 3686.2 | 86.9 |
| + epitope prediction | -1.9 | 14.4 |

##### protease

| | $elpd_{diff}$ | $se_{diff}$ |
|---|---|---|
| logistic regression (baseline) | 0.0 | 0.0 |
| + horseshoe prior | 27.7 | 13.4 |
| + phylogeny | 297.7 | 24.5 |
| + epitope prediction | 5.1 | 4.5 |

##### reverse transcriptase

| | $elpd_{diff}$ | $se_{diff}$ |
|---|---|---|
| logistic regression (baseline) | 0.0 | 0.0 |
| + horseshoe prior | 198.9 | 27.3 |
| + phylogeny | 1668.9 | 56.3 |
| + epitope prediction | 1.7 | 9.4 |
